# Crystal structures of full length DENV4 NS2B-NS3 reveal the dynamic interaction between NS2B and NS3

**DOI:** 10.1101/2020.01.27.907089

**Authors:** Wint Wint Phoo, Abbas El Sahili, ZhenZhen Zhang, Ming Wei Chen, Chong Wai Liew, Julien Lescar, Subhash G. Vasudevan, Dahai Luo

## Abstract

Flavivirus is a genus of emerging and re-emerging arboviruses which include many significant human pathogens. Non-structural protein 3 (NS3), a multifunctional protein with N-terminal protease and C-terminal helicase, is essential in viral replication. The NS3 protease together with NS2B cofactor is an attractive antiviral target. A construct with an artificial glycine linker connecting the NS2B cofactor and NS3 protease has been used for structural, biochemical and drug-screening studies. The effect of this linker on dynamics and enzymatic activity of the protease was studied by several biochemical and NMR methods but the findings remained inconclusive. Here, we designed constructs of NS2B cofactor joined to full length DENV4 NS3 in three different manners, namely **b**NS2B_47_NS3 (**b**ivalent), **e**NS2B_47_NS3(**e**nzymatically cleavable) and **g**NS2B_47_NS3 (**g**lycine-rich G4SG4 linker). We report the first crystal structures of linked and unlinked full-length NS2B-NS3 enzyme in its free state and also in complex with Bovine Pancreatic Trypsin Inhibitor (BPTI). These structures demonstrate that the NS2B-NS3 protease mainly adopts a closed conformation. BPTI binding is not essential to but favors the closed conformation by interacting with both NS2B and NS3. The artificial linker between NS2B and NS3 tends to induce the open conformation and interfere with the protease activity. This negative impact on the enzyme structure and function is restricted to the protease domain as the ATPase activities of these constructs are not affected.

## Introduction

Flaviviruses include many significant human pathogens such as dengue virus (DENV), West Nile virus (WNV) and recently re-emerging Zika virus (ZIKV). Recent outbreak of ZIKV infections in America has caused global health concern since the infections were linked to neuropathic Guillain-Barré syndrome in adults and microcephaly in infants [1–3]. DENV, has been emerging in the past decade and is a global healthcare burden. The emergence of pandemic DENV and epidemic ZIKV infections in the past years due to globalisation and urbanisation call for countermeasures such as the development of potent antivirals against these infections.

Flaviviruses are enveloped viruses which contain a single-stranded positive-sense RNA genome of about 11 kb, with 3’ and 5’ untranslated regions (UTR) [4]. The genome encodes a poly-protein precursor which is cleaved into three structural proteins and seven non-structural proteins by host and viral proteases [4, 5]. Non-structural protein 3 (NS3) plays essential roles in viral replication and polyprotein processing and is an attractive anti-viral target [6]. The N-terminal domain of NS3 (residues 1-168) is a serine protease responsible for cleavage of polyprotein precursors into mature functional proteins [7–12]. The C-terminal domain of NS3 is an NTPase/Helicase involved in viral replication and virion assembly [13–15]. Recently, several drugs targeting the Hepatitis C virus (HCV) NS3 protease have been approved by the U.S. Food & Drug Administration (FDA) [16]. However, no NS3 inhibitor for DENV has advanced to clinical trials [7, 17, 18].

The N-terminal protease contains a catalytic triad formed by residues Ser-135, His-51, and Asp-75 [19] and requires NS2B protein as cofactor for endoplasmic reticulum (ER) membrane anchorage, proper folding, and protease activity [9, 10, 19]. Soluble and catalytically active recombinant NS2B_47_-G_4_SG_4_-NS3 protease (heraftercalled gNS2B_47_NS3 Pro) was designed by tethering central NS2B cofactor to NS3 protease by a flexible artificial glycine linker [20]. Structural studies of the flavivirus NS3 protease have been done utilizing this construct design except for the recent ZIKV protease studies [12, 21–24]. These studies using conventional glycine-linked constructs demonstrated that the NS2B N-terminus contributes to the folding of protease by inserting a β-strand to the N-terminal β-barrel of protease [11, 12, 22, 25–27]. The C-terminus of NS2B is flexible and is only observed in crystal structures where the protease is bound to an inhibitor or a substrate, suggesting that it is acting as a flap closing upon substrate binding [11, 12, 22, 26]. The free protease structures with flexible NS2B C-terminus are said to adopt an “open” conformation, while the protease-inhibitor structures with NS2B contributing to substrate binding site show a “closed” conformation. Although all crystal structures of free gNS2B_47_NS3 protease are reported to adopt the open conformation, NMR studies have shown that in solution, gNS2B_47_NS3 protease interonverts between the open and closed conformations even in the absence of an inhibitor [28–32]. These studies also showed that when NS2B and NS3 are separate polypeptides, the NS2B-NS3 protease complex is mainly in the closed conformation without the substrate [29, 32].

Structural studies on ZIKV NS2B_47_NS3 protease shed new light on this unsolved issue. Zhang et al have reported a crystal structure of unlinked ZIKV protease (bZiPro) in closed conformation without an inhibitor [33]. The biochemical studies of ZIKV NS2B-NS3 protease constructs with glycine linker (gZiPro), NS2B-NS3 enzymatic cleavage site linker (eZiPro) and bivalent unlinked NS2B NS3 protease (bZiPro) have revealed that the flexible glycine linker interferes with the protease activities resulting in lower k_cat_ [24]. Shannon et al posited that reduced product release could be the possible mechanism behind the lower activity of glycine-linked constructs based on studies carried with the DENV2 bivalent co-expressed NS2B NS3 protease (bNS2B_47_NS3 Pro) and glycine-linked NS2B NS3 protease (gNS2B_47_NS3 Pro) [34]. Optimising construct designs to obtain biologically relevant crystal structures is an important factor for structure-based drug discovery. The crystal structures of separate domains of DENV NS3 have been reported, as well as full-length NS3 together with an 18-residues cofactor region of NS2B (NS2B_18_NS3)[12, 25, 35–37]. Full-length gNS2B_47_NS3 from Murine valley encephalitis virus (MVEV) has also been reported to adopt an open conformation in the absence of inhibitor [38]. Although the protease and NTPase/helicase domains of full length DENV4 NS2B_18_NS3 showed similar folds to those in MVEV NS2B_47_-NS3, domain arrangements between helicase and protease were found to be different, consistent with the flexibility of the linker region between the two functional domains. Here we designed bivalent, enzymatic cleavage site linked and conventional flexible glycine linked NS2B cofactor with full length DENV4 NS3 constructs namely bNS2B_47_NS3, eNS2B_47_NS3, and gNS2B_47_NS3 similar to those in our previous studies on ZIKV protease [24]. We report three crystal structures of full length DENV4 NS2B_47_NS3 constructs, eNS2B_47_NS3 and gNS2B_47_NS3 in free form and two in complex with Bovine Pancreatic Trypsin Inhibitor (BPTI). The structural analysis suggests that the NS2B-NS3 protease has a preformed active site with NS2B cofactor wrapped around NS3 participating in substrate binding. The biochemical studies of the ATPase activities of full length NS3 demonstrate uncoupled enzymatic activities for the full length NS3 protein.

## Results

### Design and preparation of unlinked and linked full length NS2B_47_NS3 proteins

To overcome the problem of poor expression of wild type gNS2B_47_NS3 proteins, we mutated the protease at either (1) S135 to alanine (S135A) or (2) hydrophobic residues on the surface, L30 and F31 to serine (L30S-F31S) [35]. The protease activity of gNS2B_47_NS3 Pro and of gNS2B_47_NS3 Pro (L30S-F31S) are comparable indicating that L30S-F31S mutation does not interfere with the proteolytic activity of NS3 unlike the S135A mutation which completely abolished protease activity (S1 Fig). We replaced the glycine linker of gNS2B_47_NS3 with the NS2B C-terminal penta-peptide (VKTQR) resulting in endogenous enzyme cleavable NS2B-NS3 constructs - eNS2B_47_NS3 (S135A) and eNS2B_47_NS3 (L30S-F31S) (Fig 1A). We name this eNS2B_47_NS3 L30S-F31S construct as unlinked eNS2B_47_NS3 since the NS2B/NS3 cleavage site was fully cleaved by the protease resulting in heterodimers of NS2B cofactor peptide-NS3. The bivalent construct bNS2B_47_NS3 was designed by co-expressing NS2B cofactor and NS3 sequences which fold as a heterodimer, similar to bZiPro [24, 33].SDS-PAGE analysis of proteins showed that eNS2B_47_NS3 L30S-F31S undergoes complete proteolysis resulting in unlinked full length NS3 similar to ZIKV eZiPro [24] (Fig 1B). The constructs and mutations are listed in Figure 1C. All full length NS2B_47_NS3 proteins were soluble and monomeric in solution as shown by the size exclusion chromatography profiles (S2 Fig A). Internal proteolysis at NS3 was observed during and after purification for all constructs with active protease, and gNS2B_47_NS3 degraded slightly more slowly than the bNS2B_47_NS3 and eNS2B_47_NS3 (S2 Fig B).

**Fig 1.**
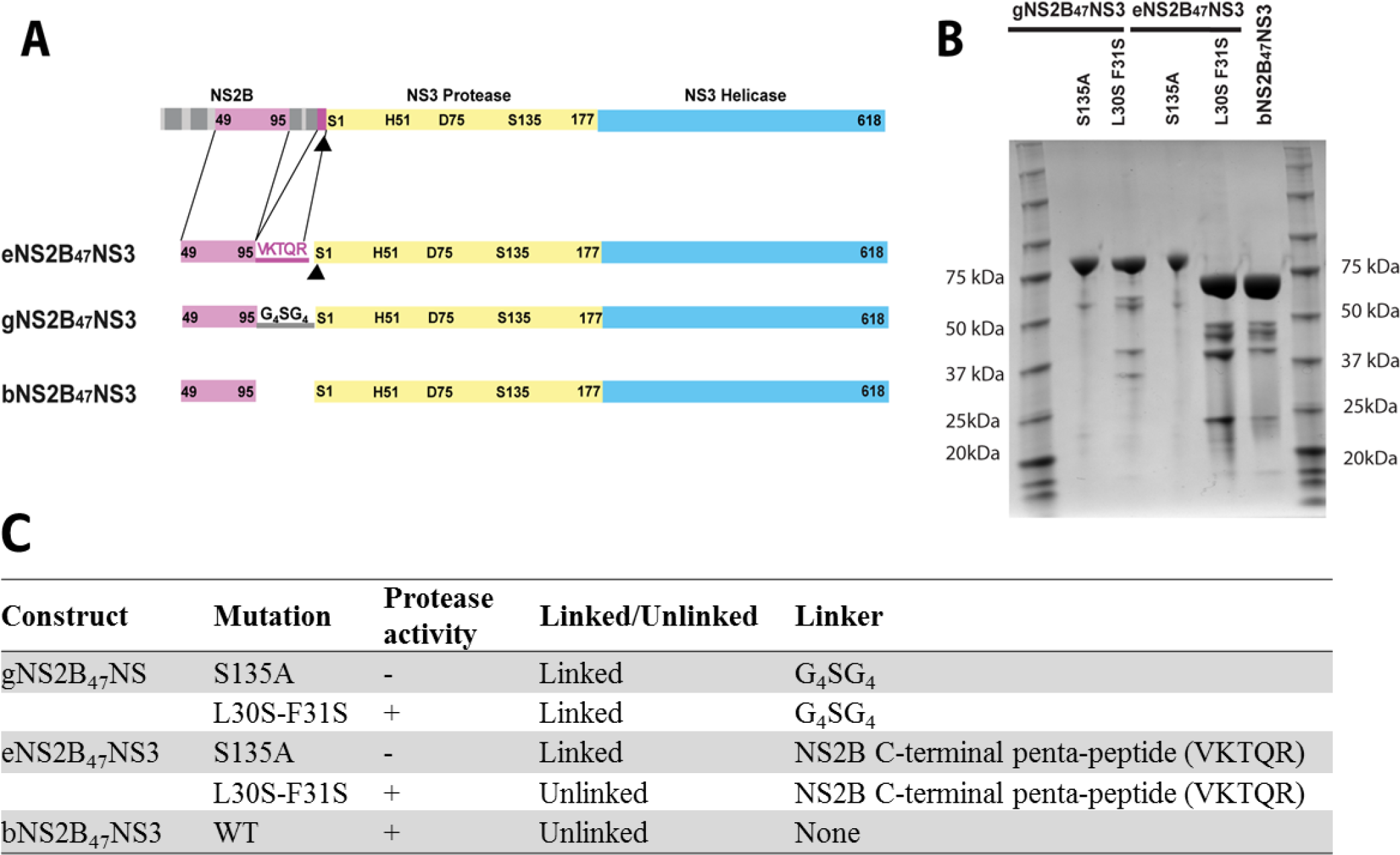
Construct design and crystal structures of DENV4 NS3. (A) Graphical representations of natural NS2B-NS3 as part of the native polyprotein, and the constructs discussed in this work. Construct-boundaries and catalytic residues are indicated. NS2B cofactor is depicted in magenta (hydrophilic region) and gray (transmembrane regions). NS3 is represented in yellow for protease domain and cyan for helicase domain. Black arrowhead indicates site of cleavage by NS3. For eNS2B_47_NS3 construct, five amino acid residues from NS2B/NS3 cleavage site is inserted between NS2B cofactor and NS3. For gNS2B_47_NS3 construct, conventional artificial flexible linker (G_4_SG_4_) is used to covalently link the NS2B cofactor and NS3. For bNS2B_47_NS3, each T7 promoter site is cloned in front of NS2B and NS3 resulting in co-expression of the cofactor NS2B and NS3. (B) SDS-PAGE analysis of purified NS3 proteins. The first and last lanes are molecular weight markers. The construct names and mutations are indicated. The eNS2B_47_NS3 L30S F31S and bNS2B_47_NS3 migrated to similar size on the gel indicating the complete proteolysis between NS2B cofactor and NS3 for the eNS2B_47_NS3 constructs. (C) The constructs are listed in the table with the types of mutation, active/inactive protease, and the type of linker between NS2B and NS3.

### Structures of full length NS2B_47_NS3

The NS2B_47_NS3 proteins were crystallised using very similar conditions (S1 Table). The crystalswere assigned into three groups that are related to the protein conformation: (1) Open conformation in which the C terminal region of the cofactor NS2B was disordered (Fig 2A); (2) -Closed conformation in which the C-terminus of NS2B loosely forms a beta hairpin (Fig 2B, 2D);(3) BPTI-bound closed conformation with similar but less dynamic NS2B C- terminus beta hairpin (Fig 2C, 2E). The correlation between the overall conformations and the unit cell dimensions is apparent from Table S1. The open conformation was only captured in gNS2B_47_NS3 (Fig 2A), which closely resembles the structure of DENV4 gNS2B_18_NS3 conformation I (PDB id: 2VBC) [35], while the remaining free enzyme structures are in closed conformation (Fig 2B, 2D). The gNS2B_47_NS3-BPTI structure and eNS2B_47_NS3-BPTI structures adopt the same conformation (Fig 1E, G). On the other hand, the bNS2B_47_NS3 protein crystals diffracted poorly to about 4 Å and as a result, we failed to find a convincing structure solution by molecular replacement. The unit cell dimensions of bNS2B_47_NS3 crystals were similar to the closed conformations of gNS2B_47_NS3 and of unlinked eNS2B_47_NS3, suggesting that the bNS2B_47_NS3 was also in a closed conformation (S1 Table).The data collection and refinement statistics are summarized in Table 1.

**Fig 2.**
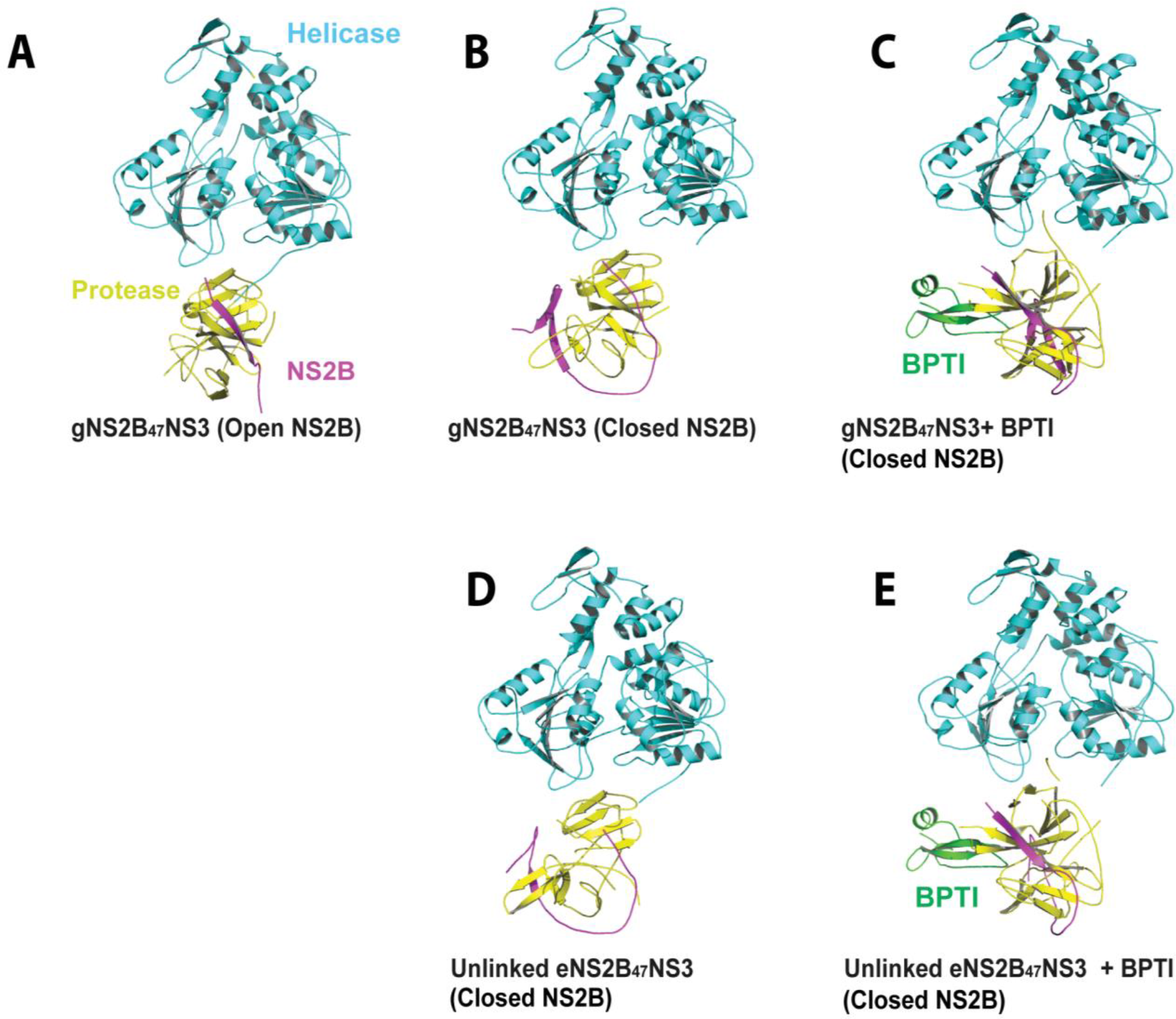
Crystal structures of NS2B_47_NS3 in apo-state and in complex with BPTI. The individual domains of NS3 and BPTI are labelled. The NS3 helicase domain is coloured in cyan, the NS3 protease in yellow and NS2B cofactor region in magenta. All five structures adopt an elongated conformation similar to previous NS2B_18_-NS3 full-length structure by Luo et al (1). The open and closed state of NS2B for each structure is stated together with the construct name. Both open NS2B and closed NS2B conformations are observed for gNS2B_47_NS3 free enzyme structures (A) and (B). (C) gNS2B_47_NS3 in complex with BPTI. (D) Unlinked eNS2B_47_NS3 structure is shown with the closed NS2B cofactor without a substrate/inhibitor. (E) Similar to gNS2B_47_NS3-BPTI structure, the NS2B C-terminus is in closed conformation for unlinked eNS2B_47_NS3-BPTI structure.

**Table 1:**
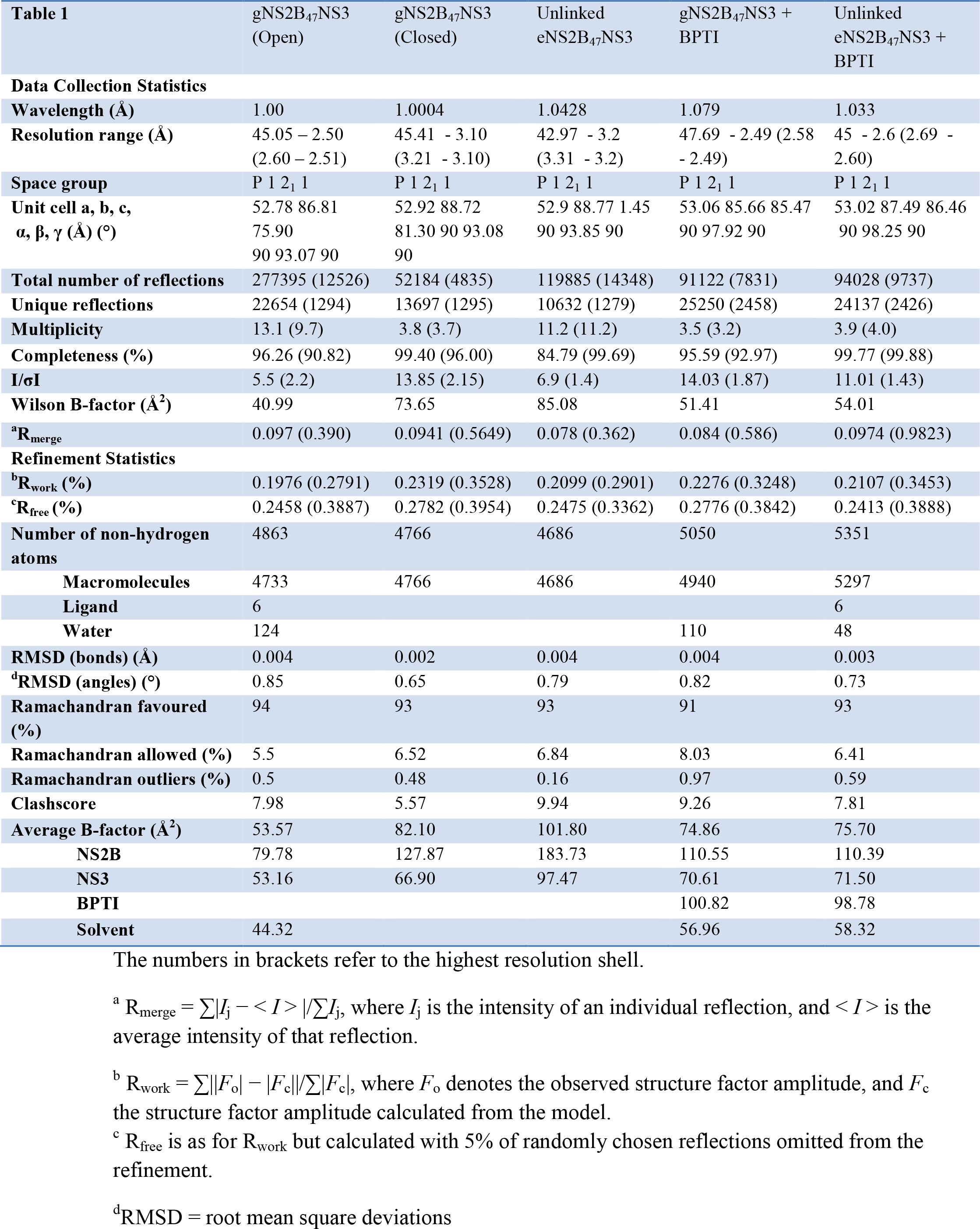
Data collection and refinement statistics

Overall, the NS2B_47_NS3 structures adopt an extended shape where the N terminal protease and the C terminal helicase domains are loosely connected through a flexible interdomain linker similar to DENV4 gNS2B_18_NS3 structures (PDB id: 2VBC, 2WHX, 2WZQ) and the MVEV gNS2B_47_NS3 (PDB id: 2WV9) (Fig 2, S3 Fig) [35, 37, 38]. In all NS2B_47_NS3 crystals, major crystal contacts are formed between the neighbouring helicase domains, which allow the protease domain to adopt multiple conformations (S4 Fig). From the open to the closed conformation, the protease domain is translated by about 0.7 Å. From the closed free enzyme conformation to the closed protease-BPTI conformation, the protease is rotated by an angle of 52.9°. Although of low resolution, the solvent shells around the protease domain are discernible confirming that the structure solution is correct (S5 Fig). Compared to MVEV NS2B_47_NS3 structure, when their helicase domains are superimposed, the protease domain are rotated by an angle of 177.1° and a translation of 17 Å (S3 Fig) with respect to the superimposed helicase domain.

We captured three free enzyme full length NS2B_47_NS3 structures. For the gNS2B_47_NS3 construct, the NS2B cofactor is captured in both open and closed conformations (Fig 3A, Fig 3B) while for unlinked eNS2B_47_NS3 constructs, the NS2B co-factor is captured only in closed conformation (Fig 3D, Fig 3E). This indicates that the presence of a flexible glycine linker increases the population adopting an open conformation. The RMSDs of protease domain between gNS2B_47_NS3, linked eNS2B_47_NS3, unlinked eNS2B_47_NS3 are less than 0.7 Å for 160 Cα atoms. In the free enzyme structures with closed NS2B conformation, the last 10-12 amino acids from the cofactor NS2B, the linker, and the first 20 amino acids of NS3 are flexible. In eNS2B_47_NS3 structures, the NS2B N terminal 8 residues and 3 residues are flexible for linked and unlinked structures respectively. The electron density of NS2B C-terminus for free enzyme closed NS2B conformation structures are relatively weak indicating that the C-terminus of NS2B is dynamic when the active site is not occupied (Fig. 3).

**Fig 3.**
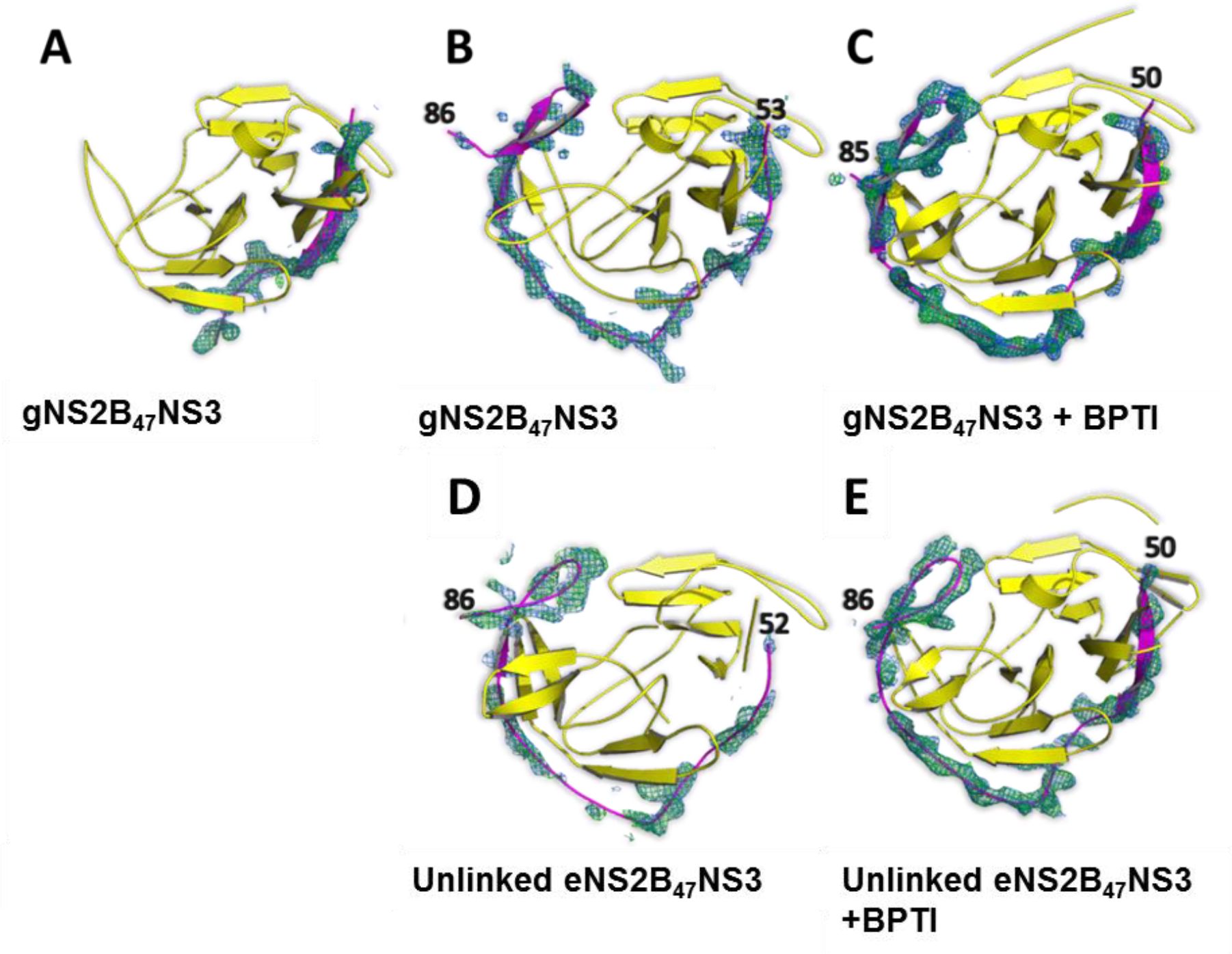
Different conformations of NS2B cofactor in full length NS3 structures. The NS2B is shown in magenta and NS3 in yellow. The 2F_o_-F_c_ map contoured at a level of 1 σ is shown in blue and F_o_-F_c_ map contoured at 3 σ is shown in green, where the NS2B was omitted in the calculation. (A, B) The protease domain of gNS2B_47_NS3 with cofactor NS2B shows that NS2B could adopt both open and closed conformations without the inhibitor. (C, E) When the protease is in complex with BPTI, NS2B cofactor is in closed conformation for both gNS2B_47_NS3 (C) and eNS2B_47_NS3 structures (F). (D) The NS2B cofactor of unlinked eNS2B_47_NS3 protease stays in a closed conformation without inhibitor. The electron density maps of NS2B in free enzyme structure is weaker than that of NS2B in protease-BPTI complex structure indicating that without substrate NS2B is dynamic.

Interestingly in the unlinked eNS2B_47_NS3 structure, the NS2B/NS3 cleavage peptide (VKTQR) is not occupying the substrate binding site, unlike in the similar ZIKV protease structure, (eZiPro) (PDB accession code 5GJ4) [24]. We also report the crystal structures of gNS2B_47_NS3 and of unlinked eNS2B_47_NS3 in complex with BPTI. The RMSD between the two NS3-BPTI structures are 0.44 Å for 618 Cα atoms indicating that mode of binding of BPTI is conserved in both gNS2B_47_NS3 and eNS2B_47_NS3. The protease domain rotates by 52.9° in the NS3-BPTI full length structure to accommodate the BPTI in the crystal. The detailed interactions between the BPTI and NS2B cofactor and NS3 protease in both structures are conserved (S6 Fig). The three NS2B_47_NS3 free enzyme structures reveal a more dynamic NS2B cofactor and NS3 protease compared to the two NS2B_47_NS3-BPTI structures indicating that substrate binding stabilises the protease (Fig 3, S7 Fig). The ATPase/helicase domains of both gNS2B_47_NS3 and eNS2B_47_NS3 are identical with RMSD of less than 0.5 Å. The overall conformation of helicase is similar to the helicase structures with no NTP or RNA bound except for the residues 461-471 as mentioned before by [38] and residues 243-253. This surface loop is in close proximity to NS2B β hairpin and to NS3 residues 66PSWAD71, and changes conformation when the BPTI binds to the protease domain. In eNS2B_47_NS3-BPTI structure, movement of the protease domain results in the P-loop moving away from these residues.

### The artificial glycine linker interferes with the protease activity of NS3

Latest studies using biochemical and NMR of flaviviral protease have shown that the flexible glycine linker affects the enzymatic and binding activities of the protease [23, 24]. To determine the effect of artificial glycine linker on the enzymatic activities of full length DENV NS3, we measured the protease activity of eNS2B_47_NS3, gNS2B_47_NS3, and bNS2B_47_NS3 using Benzoyl-Nle-Lys-Arg-Arg-Aminomethylcoumarin (Bz-NKRR-AMC) fluorescent substrate [39]. Our enzymatic assays showed that while the glycine linker does not affect the substrate apparent affinity (*K_m_*), its presence slows down the rate of catalysis (*k_cat_*) (Fig 4A). It is possible that the glycine linker introduces steric hindrance on the NS2B- NS3 conformational transitions compared to unlinked construct. Although eNS2B_47_NS3 has a slightly higher *K_m_* and lower *k*_cat_ compared to bNS2B_47_NS3, presence of NS2B/3 cleavage site does not have the similar inhibitory effect on the protease enzymatic activity as reported for eZiPro [24]. This could be due to the sub-optimal cleavage site found at NS2B/NS3 in all DENV serotypes where the P2 residue is glutamine instead of a strongly basic lysine or arginine found in other flaviviruses. The inhibition activity assays with BPTI and with small peptidic inhibitor, Benzoyl-Lys-Arg-Arg-H, shows that the half maximal inhibitory value, IC_50_, was lowest for bivalent bNS2B_47_NS3 (Fig 4B,C) and highest for gNS2B_47_NS3 indicating a slightly tighter association with the former. In addition, the thermal shift assay of these constructs shows T_m_ of gNS2B_47_NS3 is 2°C lower than that of bNS2B_47_NS3 and of eNS2B_47_NS3, further suggesting that the presence of artificial flexible linker between NS2B cofactor and NS3 may interfere with the protein stability (S2 Fig).

**Fig 4.**
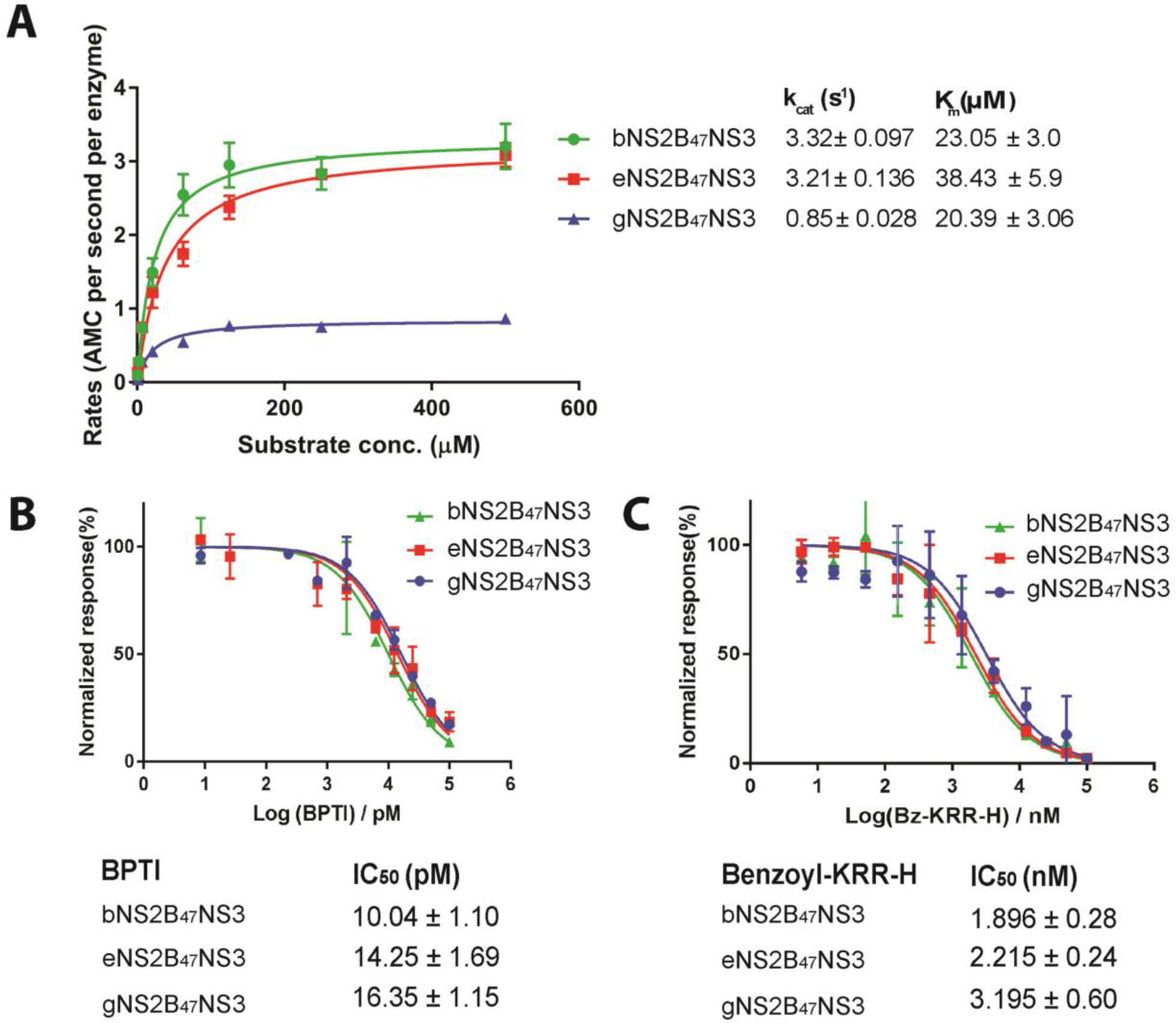
Characterisation of protease activity of NS3 full length constructs with different linkers. (A) Protease activity of bNS2B_47_NS3, eNS2B_47_NS3 L30S F31S and gD4NS2B_47_NS3 L30S F31S against Benzoyl-Nle-Lys-Arg-Arg-AMC substrate. (B,C) Half maximal inhibition efficiencies (IC_50_) of BPTI and Benzoyl-Lys-Arg-Arg-H against NS2B_47_NS3 constructs were determined. The gNS2B_47_NS3 showed lowest *k_cat_* and *K_m_.* The presence of NS2B-NS3 cleavage site does not affect the enzymatic activities of full length NS3 as seen by comparable *k_cat_*s between bNS2B_47_NS3 and eNS2B_47_NS3.

### The kinetics of ATP hydrolysis by full length NS3 constructs are similar

Next, to determine the effect of linkers on the NTPase activities of NS3, we carried out the NADH coupled ATPase assay for g-, e-, bNS2B_47_NS3 full length constructs. These constructs show similar ATPase activity demonstrating that the different linkers between NS2B and NS3 protease do not interfere with NTPase activity of the helicase. The helicase activity of NS3 requires the energy provided by ATP hydrolysis. The NTP binding site of helicase is situated right on top of the protease domain while the RNA binding groove of the helicase domain is spatially separated from the protease domain. Therefore, these different linker constructs are unlikely to have an effect on the helicase activity if the ATPase activity is unaffected by the presence of different linkers between NS2B and NS3. To test if the binding of substrate to the protease domain affect the NTPase activity of helicase domain, we measured the ATPase activity of bNS2B_47_NS3 in the presence and absence of BPTI. The rate of ATP hydrolysis remains unchanged when BPTI is bound to the protease domain, demonstrating that the substrate binding on protease domain does not affect the ATPase activity of helicase domain (Fig 5B). The ATPase activity of DENV4 NS3 helicase was measured in the presence and absence of BPTI as the control (Fig 5B). Both *k_cat_* and *K_m_* of ATP hydrolysis of bNS2B_47_NS3 is slightly slower compared to those of helicase alone, and hence the catalytic efficiencies of both enzymes are similar (Fig 5B).

**Fig 5.**
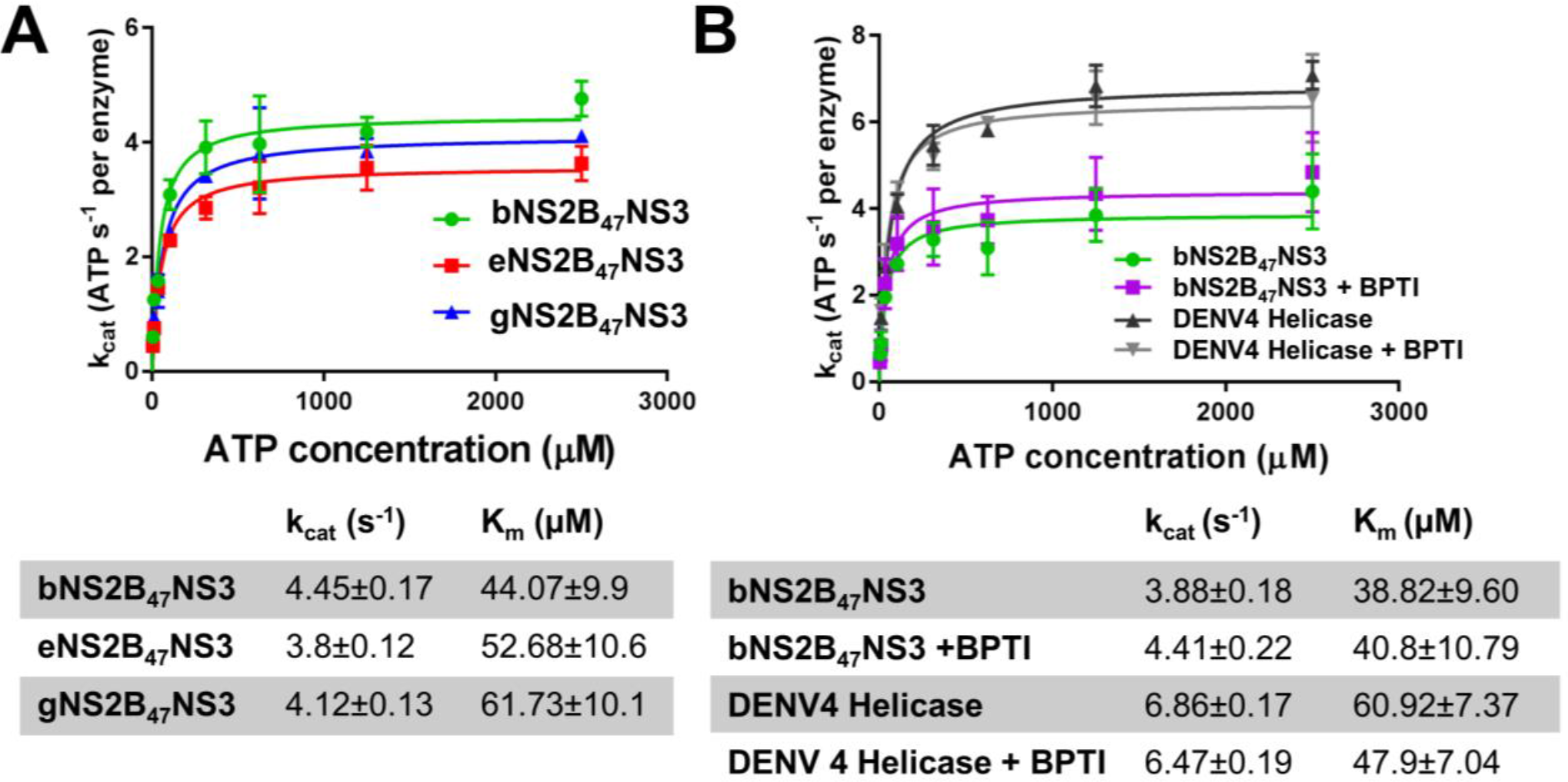
The ATP hydrolysis activity of NS2B_47_NS3 constructs were compared along with Helicase. (A) Rate of ATP hydrolysis of bNS2B_47_NS3, eNS2B_47_NS3 and gNS2B_47_NS3. The Michaelis-menten parameters are stated below the curves. The presence of artificial glycine linker and of NS2BNS3 cleavage junction slightly lowers the ATP hydrolysis by increasing K_m_. (B) ATP hydrolysis of bNS2B_47_NS3 with or without BPTI shows that presence of inhibitor does not interfere with ATP hydrolysis. The DENV4 helicase was used as positive control.

## Discussion

Due to absence of NS2B/NS3 crystal structures in closed conformation without substrate or inhibitor, NS2B was proposed to convert from open and closed conformations upon substrate binding [11, 12, 25–27]. Early NMR studies of glycine linked DENV and WNV proteases showed crowded cross peaks due to conformational exchanges [28, 30]. The use of unlinked constructs in the followed-up NMR studies has improved the spectral quality and backbone assignment [29, 32]. The unlinked DENV protease constructs are obtained by 1) replacement of glycine linker with NS2B/NS3 cleavage site (EVKKQR) similar to eNS2B_47_NS3 and 2) by co-expression of NS2B cofactor peptide and NS3 protease similar to bNS2B_47_NS3 [29, 32]. The NMR studies of these unlinked DENV proteases confirmed that the NS2B cofactor is predominantly in a closed conformation. Likewise, NMR studies for similar ZIKV protease, eZiPro, bZiPro, and gZiPro, also showed that the unlinked protease is in a closed conformation [23, 24, 40]. The presence of glycine linker between NS2B and NS3 shifts the population towards open NS2B conformation, leading to crowded peaks in NMR spectra, whereas for unlinked NS2B-NS3 protease, well-resolved spectra are obtained due to the dominant closed NS2B conformation [29, 32, 40]. Here, we report a series of crystal structures of DENV4 NS2B_47_NS3 protease-ATPase/helicase which were designed in different formats and captured as free enzyme and inhibitor-bound complexes. These structures for the first time clearly confirm that both gNS2B_47_NS3 and unlinked eNS2B_47_NS3 could adopt the closed NS2B conformation in the absence of any substrate or inhibitor. These results therefore demonstrate that NS2B_47_NS3 protease has a preformed ligand binding site which becomes further stabilized upon substrate binding. For unlinked eNS2B_47_NS3, the NS2B/NS3 cleavage site pentapeptide (VKTQR) is not found at the active site, in contrast to the otherwise comparable protease structure from ZIKV [24]. All the structures reported here are crystallised under similar crystallization conditionsand the major crystal contacts are formed by the helicase domain (S6 Fig). This implies that these constructs could be further engineered to study the structural properties of NS2B-NS3 protease. The NS2B/3 cleavage pentapeptide of eNS2B_47_NS3 could be replaced by othercleavage sites present in the viral polyprotein. Determination of the crystal structures of the above mentioned constructs could be useful in understanding how different polyprotein cleavage sites bind to the NS2B-NS3 protease. Moreover, the binding loop of BPTI could be mutated as reported by Lin et al and subsequently co-crystallised with eNS2B_47_NS3 as a scaffold to understand the prime site interactions between the inhibitor and the protease [41].

The protease activity assays of different constructs show that bNS2B_47_NS3 and unlinked eNS2B_47_NS3 have comparable *k_cat_* while gNS2B_47_NS3 displays the lowest *k_cat_* (Fig 4A). The ATPase activity of these constructs are similar. This indicates that the lower *k_cat_* in the protease activity observed for gNS2B_47_NS3 is unlikely due to other factors, such as small differences in enzyme concentrations or in protein stability (Figure 5A). The flexible glycine linker might introduce steric hindrance on NS2B dynamics and therefore lower the *k_cat_*. Both BPTI and peptidomimetic inhibitor, Bz-KRR-H, inhibit the protease activity of all three constructs with similar range of affinity (Fig 4B,C) indicating that the flexible linker is not interfering with inhibitor or substrate binding. Hence, it is plausible that the dynamics of NS2B and NS3 are involved at the post-catalytic/product-release stage rather than simply during substrate binding. This is in accordance with the single molecule enzymatic studies performed by Shannon et al, where the *k_cat_* of the enzyme was affected rather than *K_m_* when the NS2B-NS3 interactions were disrupted [34]. In agreement with the crystal structure, the enzymatic activities of eNS2B_47_NS3 are similar to bNS2B_47_NS3 again in contrast to that of eZiPro and bZiPro [24]. These results suggest that DENV NS2B NS3 cleavage site is released from the active site upon cleavage, whereas for ZIKV, it remains bound at the active site. It is possible that different flavivirus are employing the different polyprotein cleavage site and specificity to regulate the protease activity of NS3 *in vivo*. From our structures, we propose that NS2B/NS3 protease mainly stays in closed conformation regardless of the presence of a substrate. During polyprotein processing, NS2B is anchored to ER membrane by N and C-terminal hydrophobic regions [9, 42]. The complete dissociation of NS2B C-terminus from NS3 protease would not be favourable spatially due to the NS2B membrane anchorage, whereas stable tight association of whole NS2B cofactor to NS3 will place the active site of NS3 close to the membrane, shielding it from substrate binding or interfering with substrate release Therefore, a rather loosely associated NS2B-cofactor appears to be the optimal conformation for NS2B-NS3 *in vivo*.

In conclusion, we provide crystallographic evidence that the NS2B cofactor loosely assumes closed conformation around NS3 protease in the full length NS3 in the absence of substrate. In contrast to the unlinked ZIKV protease, eZiPro, the substrate pocket of eNS2B_47_NS3 is not occupied and therefore may be useful for co-crystallisation with inhibitors for antiviral drug discovery. Due to slightly better protease activities, bNS2B_47_NS3 and eNS2B_47_NS3 appear to be better suited for more sensitive high-throughput screening of potential drugs.

## Materials and methods

### Plasmid preparation

The bacterial expression plasmid containing wild type NS3 linked to cofactor NS2B residues 49—95 was generated by site directed mutagenesis method by inserting NS2B 68-96 to the gNS2B_18_NS3 construct from Luo et al [35]. The eNS2B_47_NS3 construct was generated by replacing the glycine linker with residues 126-130 of NS2B C-terminus which is the enzymatic cleavage site of NS2B/NS3. The eNS2B_47_NS3 L30S F31S and gNS2B_47_NS3 L30S F31S are mutated from eNS2B_47_NS3 WT and gNS2B_47_NS3 WT by site directed mutagenesis. The bivalent full length construct (bNS2B_47_NS3) was synthesized by biobasic.

### Expression and purification

The plasmids containing bNS2B_47_NS3, gNS2B_47_NS3, eNS2B_47_NS3 or mutants were transformed into *Escherichia coli* BL21(T1R). The transformants were grown in Luria Broth (LB) medium supplemented with suitable antibiotics (ampicillin (100 mg/L) or kanamycin (50 mg/L) and chloramphenicol (37 mg/L)), 40-50 mM Potassium Phosphate buffer pH 7.4 and 2.5% glycerol at 37°C until OD_600_ of 0.8 was reached. The culture was cooled to 18°C, subsequently induced with 1mM Isopropyl β-D-1-thiogalactopyranoside, and the proteins were overexpressed overnight at 18°C shaking at 200 rpm. Cells were harvested after 15 hours by centrifugation at 5000 rpm for 20 minutes at 4°C. Cells were resuspended in lysis buffer (25 mM 4-(2-hydroxyethyl)-1-piperazineethanesulfonic acid (HEPES), pH 7.5, 500 mM Sodium chloride (NaCl), 5 mM β-Mercaptoethanol (β-ME), 5% glycerol, 10 mM imidazole). Cells were lysed by passing though NIRO SOAVI PANDA HIGH PRESSURE HOMOGENIZER at pressure 700-900 bars. The soluble fraction was separated by centrifugation of the lysate at 40000 RPM for 40 minutes. The soluble proteins were purified by metal affinity chromatography using Ni-NTA beads (Thermofisher). The N terminal Histidine-tag was cleaved by Tobacco Etch Virus (TEV) protease while the eluted fraction was dialyzed against Size exclusion chromatography (SEC) buffer (25mM HEPES, pH 7.5, 150 mM NaCl, 2 mM DTT, 5% glycerol) overnight at 4°C. The His-tag cleaved proteins were further purified by running through HiTrap Heparin HP 5 ml column (GE Healthcare) and were finally polished with size exclusion chromatography using HiLoad 16/600 Superdex 200 (GE Healthcare).

### Crystallization, data collection and refinement

Crystals were grown by mixing 1 µL of proteins at a concentration of 8.5 mg/ml with 1 µL of precipitant by hanging drop vapour diffusion method (S1 Table). Cluster of thin plate crystals grew after 2 days of incubation at 20 °C. Crystals are separated into single plates, transferred to cryoprotected reservoir solution with 20% glycerol and cooled down to 100 K in liquid nitrogen before mounting.

Diffraction intensities were recorded on PILATUS 2M-F detector at PXIII beamline at the Swiss Light Source, Paul Scherrer Institut, Villigen, Switzerland and on ADSC Quantum 210r Detector at MX1 beamline at Australian Synchrotron. Diffraction intensities were integrated using iMOSFLM or XDS [43–45]. Scaling and merging of the intensities were done using POINTLESS and AIMLESS from CCP4 suite [46–49].Data collection statistics are summarized in (Table 1). For gNS2B_47_NS3 and eNS2B_47_NS3 unlinked datasets, the multiplicity was higher due to the smaller oscillation of the Φ.

The solution for gNS2B_47_NS3 with BPTI was solved using PHASER MR (CCP4 suite) using 2VBC as search model [37]. The solutions for full length NS3 (gNS2B_47_NS3 and eNS2B_47_NS3) were solved by using PHASER MR (CCP4 suite) using gNS2B_47_NS3 free enzyme structure as search model. The dataset for unlinked eNS2B_47_NS3 has ice rings and therefore the diffractions spots at the resolution shells around 3.4 Å were removed to reduce the noise. This has resulted in lowered completeness of the overall dataset. The structure solutions were subject to rounds of refinement using Phenix.refine program and manual refinement using WinCoot[50–54]. Rotational and translational movements of domains were carried out using DynDom (CCP4 suites) and Superpose (CCP4 suites)[55, 56]. Figs were generated using Pymol and electron density maps were generated using FFT (CCP4 suites)[57, 58]

### Protease activity assay

The protease activity assays were carried out using 7-amino-4-methylcoumarin (AMC) fluorophore, Benzyonyl-Nle-Lys-Arg-Arg-AMC (Peptide Institute, Japan) modified from [39]. The Bz-NKRR-AMC substrate with starting concentration of 300 µM was serially diluted in assay buffer (20 mM Tris hydrochloric acid, pH 8.5, 10% glycerol, 0.01% Triton X-100, 2 mM DTT) and added to Corning® 96 Well black plates with 3 nM protein in same buffer. Assays were carried out as duplicates or triplicates at 37°C. The rate of AMC released was monitored at Synergy™ HTX Multi-Mode Microplate Reader at excitation wavelength 380 nm and emission wavelength 460 nm over 5-10 minutes at 1 minute intervals. To determine the amount of AMC released, standard AMC curve was plotted with over different concentrations of AMC (data not shown). Initial velocities were calculated using linear regression function using GraphPad Prism version 5.0 for Windows. The relative fluorescence units (RFU) were converted to amount of AMC using the standard curve. Data were analysed and plotted using Michalis-Menten equation with GraphPad Prism version 5.00 for Windows (GraphPad Software, San Diego, California, USA).

### Protease inhibition assay

The protease inhibition assays were carried out using the same substrate used in enzymatic assay at 30 µM concentrations. The inhibitors of different concentrations were added to the wells with 3 nM of proteins and were incubated for 30 minutes at room temperature. The reaction was initiated by addition of 30 µM substrate and initial velocities were measured at 1 minute intervals at 37°C for 10 minutes. Data were analysed using function Log inhibitor vs normalized response function in GraphPad Prism.

### ATPase assay

ATPase activity assay was carried out based on Kiianitsa et al[59]. 50 nM of enzymes were incubated in assay buffer (25 mM MOPS pH 7.4, 150 mM potassium chloride, 2 mM DTT, 0.01% Triton X-100) with 50 µM of BPTI for an hour in Corning® 96 Well clear plates. NADH mixture (NADH 1mM, Phosphoenol pyruvate 2.5mM Pyruvate Kinase 500 U/ml and lactic dehydrogenase 100 U/ml in ATPase assay buffer) was added to reaction and incubated for 30 minutes more. Reaction was started by addition of various ATP concentrations. Depletion of NADH was measured by change in absorbance at 340 nm and was plotted against time using Cytation 3 Mulitmode plate reader (BioTek). After determining the path length, molar extinction coefficient for the given path length (K_path_) was calculated. Initial velocities were calculated using linear regression function using GraphPad Software version 5.0 for Windows. Data were plotted using Michaelis-Menten equation in GraphPad Prism.

### Thermal shift assays

The Thermofluor assay was carried out as described previously [60]. The samples contained 10 µM protein and 5x SYPRO Orange dye in buffer containing 20 mM HEPES pH 7.5, 150 mM NaCl, 2 mM DTT and 5% glycerol. The samples were subject to temperature increments of 1°C from 20°C to 95°C over 20 minutes using real-time PCR machine Bio-Rad CFX96. The fluorescence intensities were recorded and analysed using GraphPad Prism. The melting curves were generated using Boltzmann-sigmoidal function.

## Acknowledgment

We thank scientists from Australian Light Source MX beam-line and Swiss Light Source PX beam-line for their help with diffraction data collection. This work was supported by (1) the start-up grant to DL lab from Lee Kong Chian School of Medicine, Nanyang Technological University, (2) Ministry of Education grant MOE2016-T2-2-097 to DL lab, (3) National Medical Research Council grant CBRG14May051 to JL, (4) National Medical Research Council grant NMRC/CBRG/0103/2016 to SV lab, (5) National Research Foundation grant NRF2016NRF-CRP001-063. Ms. Wint Wint Phoo is supported by Nanyang Research Scholarship, Nanyang Technological University.

## Supporting information

**S1 Fig.**
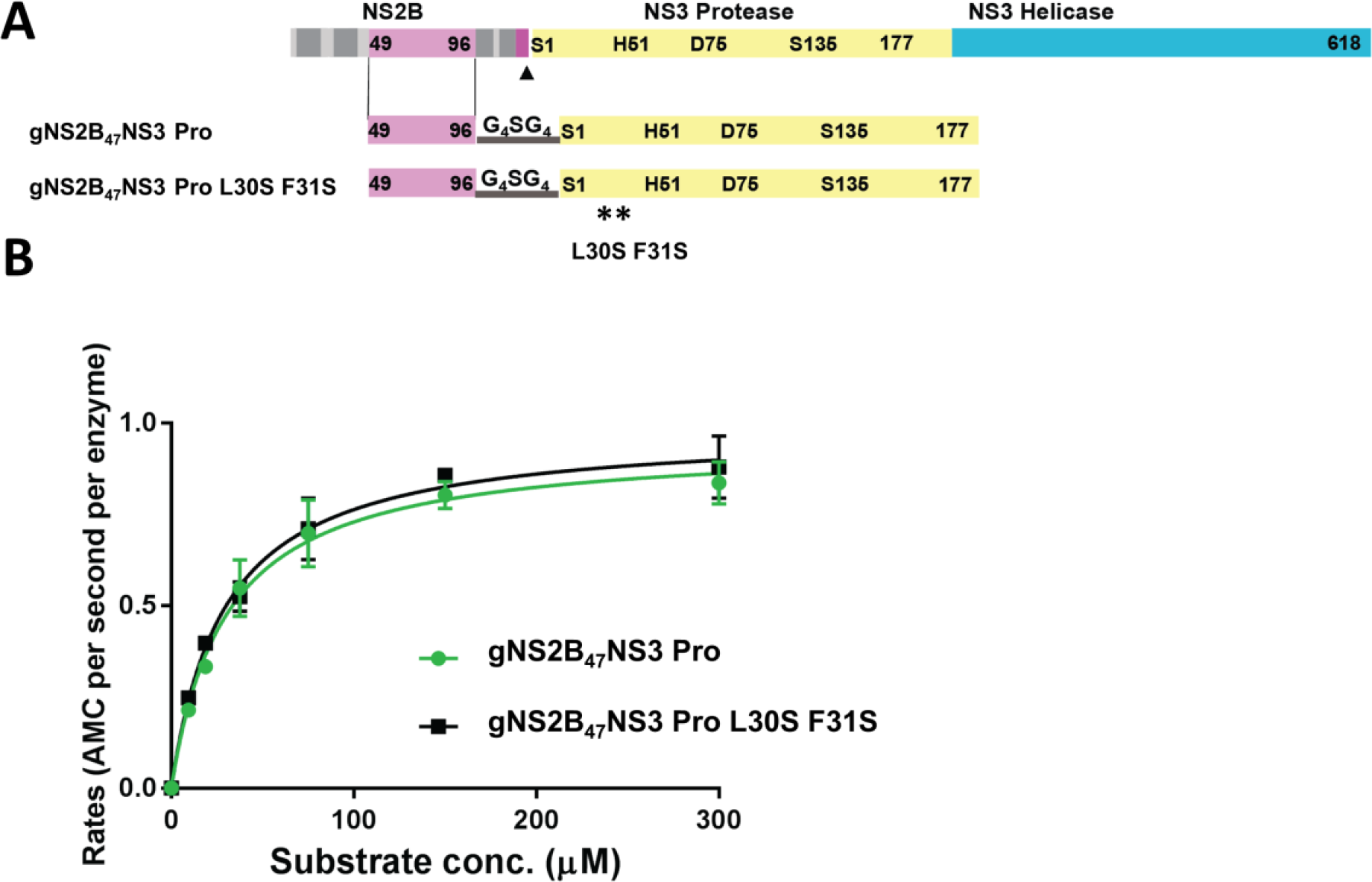
The mutations L30S F31S do not interfere with protease activity. (A) Graphical representation of NS2B NS3 gene, construct design and mutations. NS2B in magenta, and NS3 protease in yellow and NS3 helicase in cyan. Protease and helicase are labelled and boundary residues are numbered. The catalytic residues, as well as mutated residues are labelled and numbered. (B) Enzymatic activity of gNS2B_47_NS3 Pro and gNS2B_47_NS3 Pro L30S F31S. Both enzymes have similar *k_cat_* and *K_m_* indicating that L30S F31S mutations do not interfere with enzymatic activity.

**S2 Fig.**
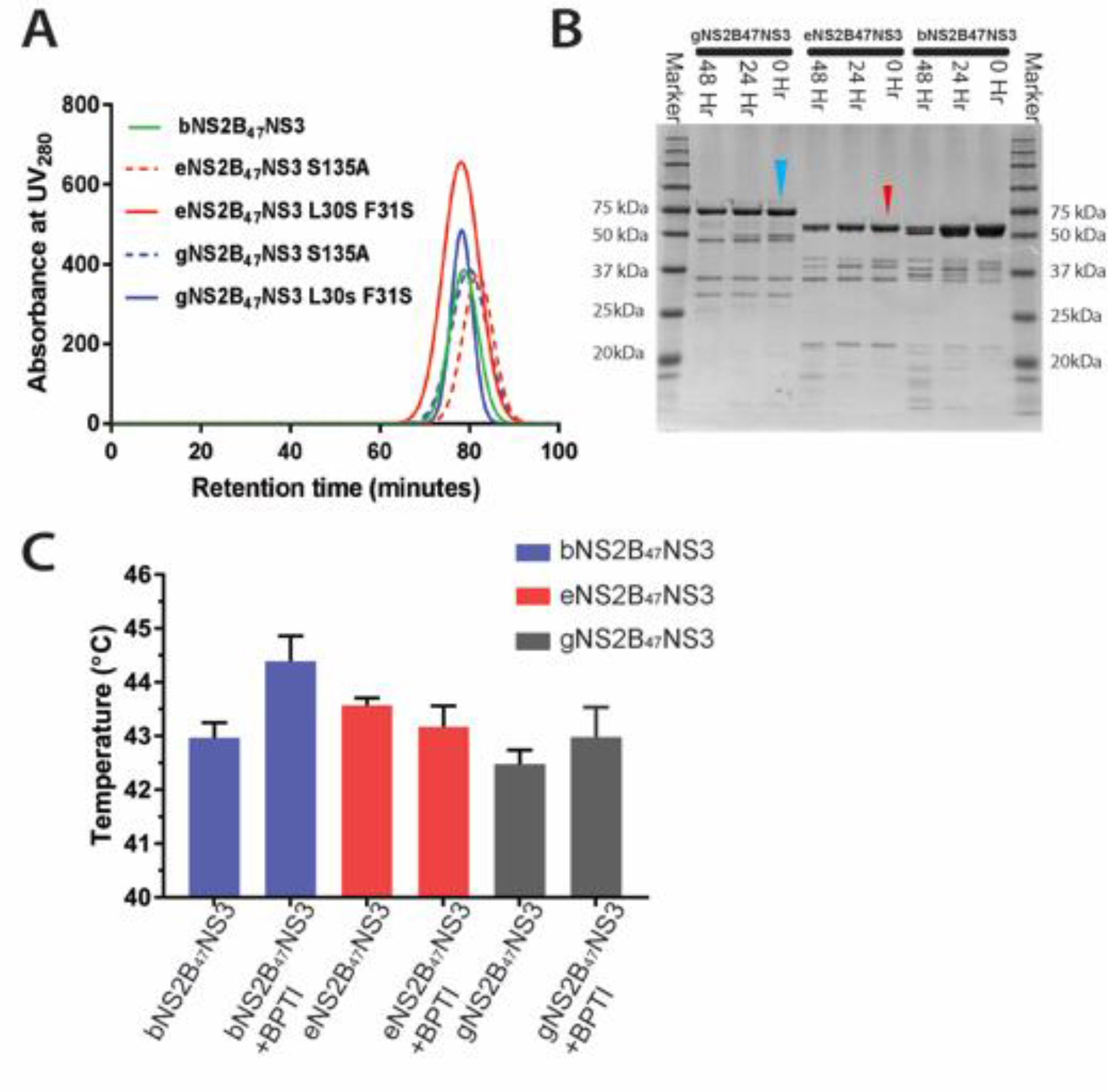
Purification of gNS2B_47_NS3, eNS2B_47_NS3 and bNS2B_47_NS3 showed monomeric proteins. (A) SEC chromatography profile of full length proteins showing that full length NS3 is monomeric. (B) SDS-PAGE analysis of full length NS3 auto proteolysis over 0hour, 24hour and 48 hours The gNS2B_47_NS3 and eNS2B_47_NS3 are indicated with blue and red arrows respectively. (C) Melting temperatures of bNS2B_47_NS3, eNS2B_47_NS3 and gNS2B_47_NS3 with/without BPTI. The gNS2B_47_NS3 has the lowest T_m_ indicating that gNS2B_47_NS3 has lower stability compared to eNS2B_47_NS3 and bNS2B_47_NS3.

**S3 Fig.**
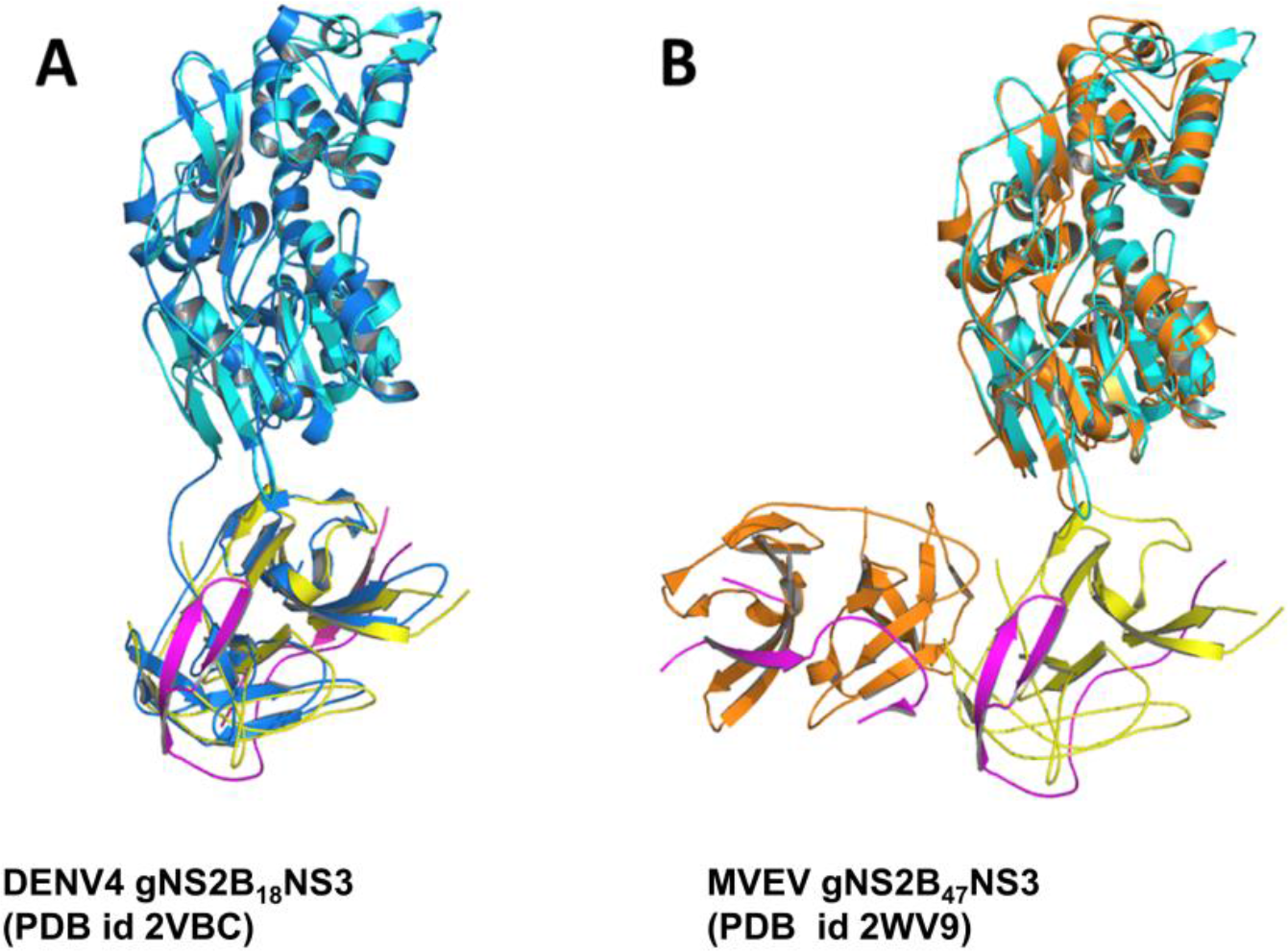
Comparison between overall conformations of NS2B_47_NS3 with previous structures. In both (A) and (B), gNS2B_47_NS3 free enzyme structure in closed conformation was shown in cyan for helicase domain, yellow for protease and magenta for NS2B (A) Superposition of current full length NS2B_47_NS3 with DENV4 full length NS3 structure with 18 residues from NS2B cofactor which is shown in blue for NS3 and magenta for NS2B (Luo et al PDB id 2VBC) (B) Superposition of current full length NS2B_47_NS3 structure with MVEV gNS2B_47_NS3 structure. The MVEV NS3 is shown in orange and NS2B in magenta.

**S4 Fig.**
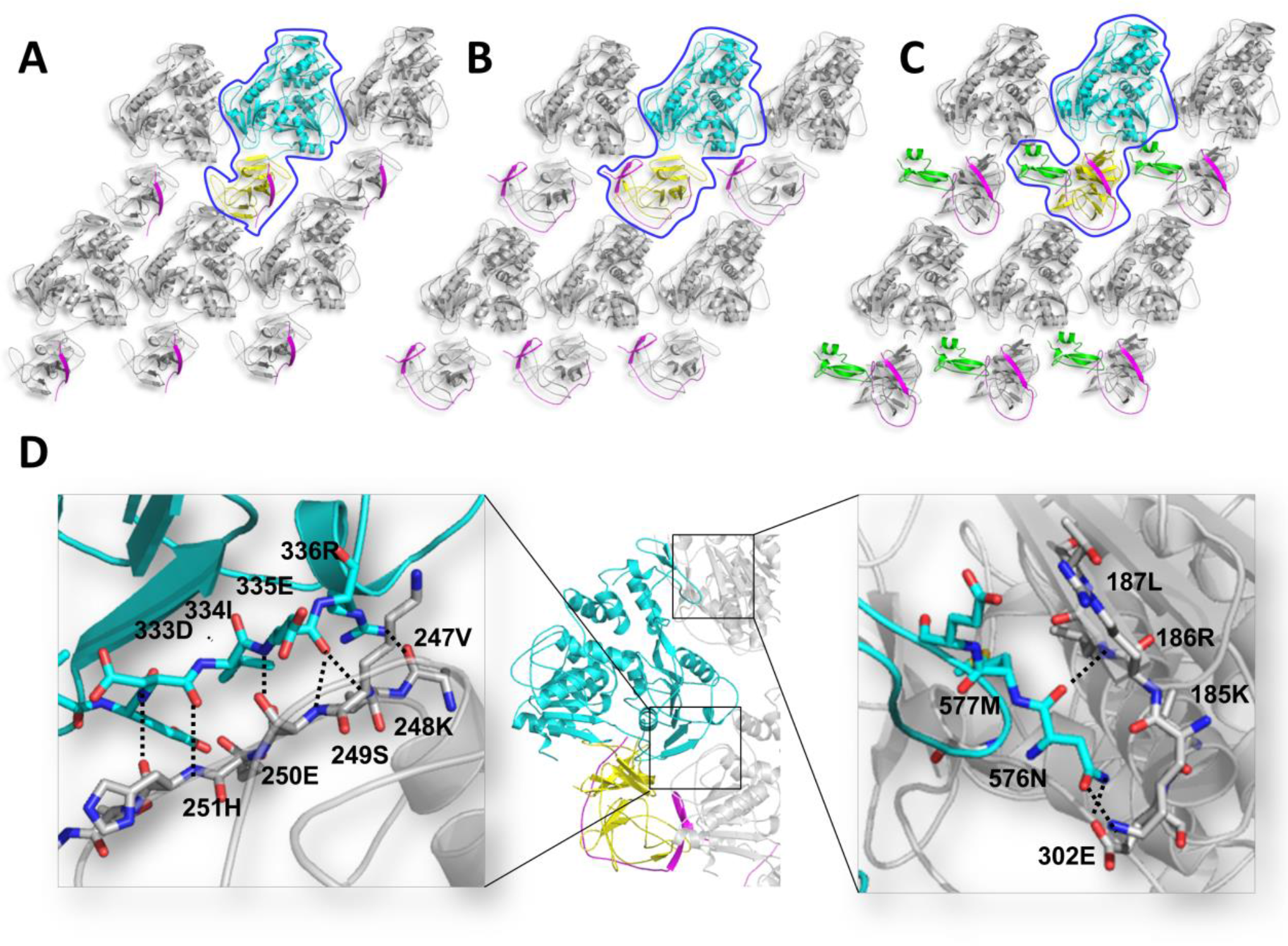
Major crystal contacts in full length NS3 structures are formed by the helicase domain. Here we display the three conformations of gNS2B_47_NS3 full length structures (A) open NS2B conformation, (B) closed NS2B conformation, (C) enzyme in complex with BPTI along with its symmetry mates. NS3 helicase domain is shown in cyan and NS3 protease in yellow. The surrounding symmetry mates are shown in grey. The NS2B is shown magenta. (D) Detailed interactions of major crystal contacts. The residues that are interacting with the symmetry mates are presented with residue number. The protease domain does not interact with neighbouring molecules giving it the conformational freedom to adopt different orientations.

**S5 Fig.**
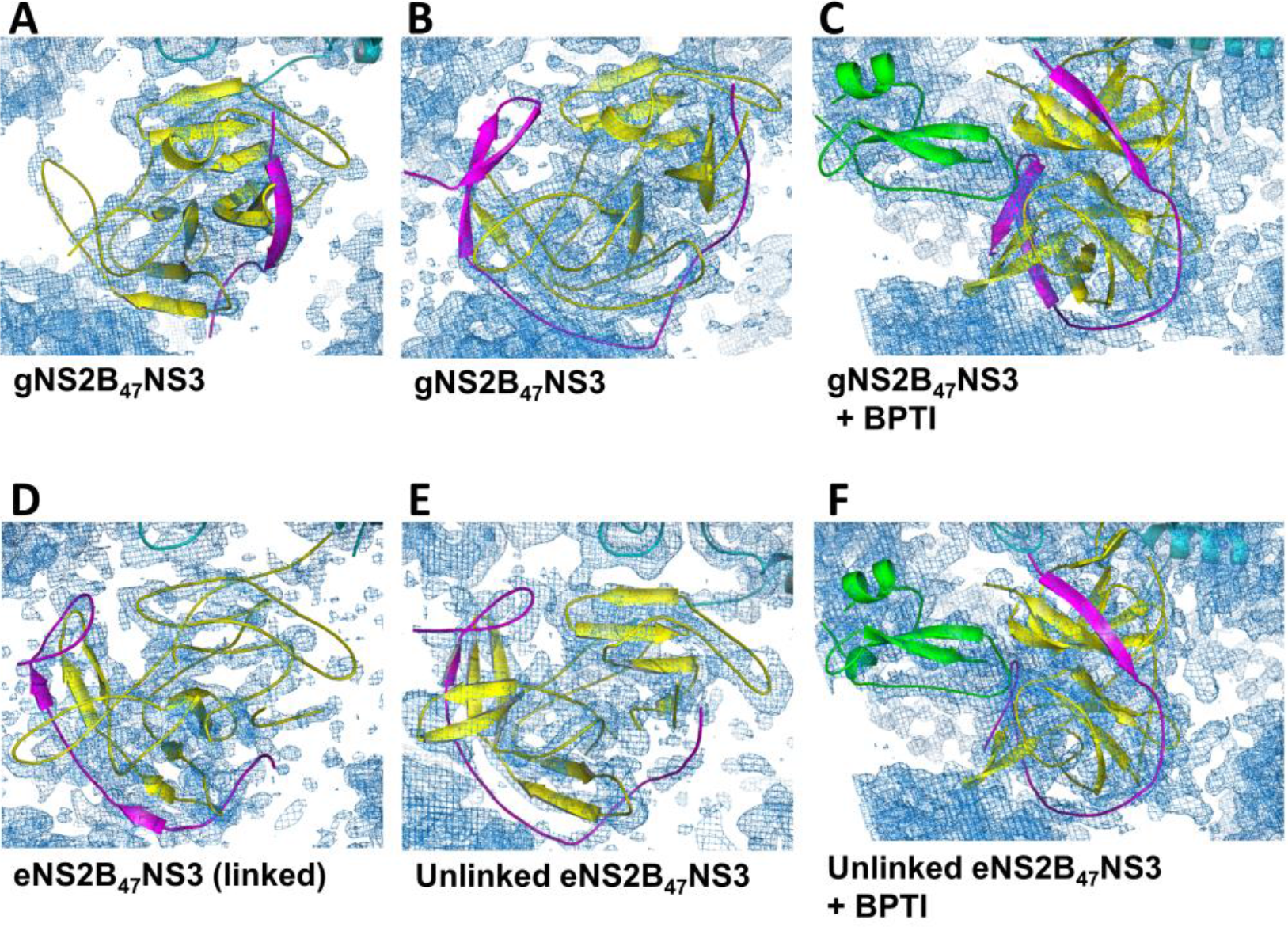
The 2mF_o_-F_c_ maps of NS2B_47_NS3 structures display solvent shell around the electron density of protein. This indicates that the structure solutions for the protease domain are correct and refined. The NS2B is colored in magenta, NS3 protease in yellow and BPTI in green.

**S6 Fig.**
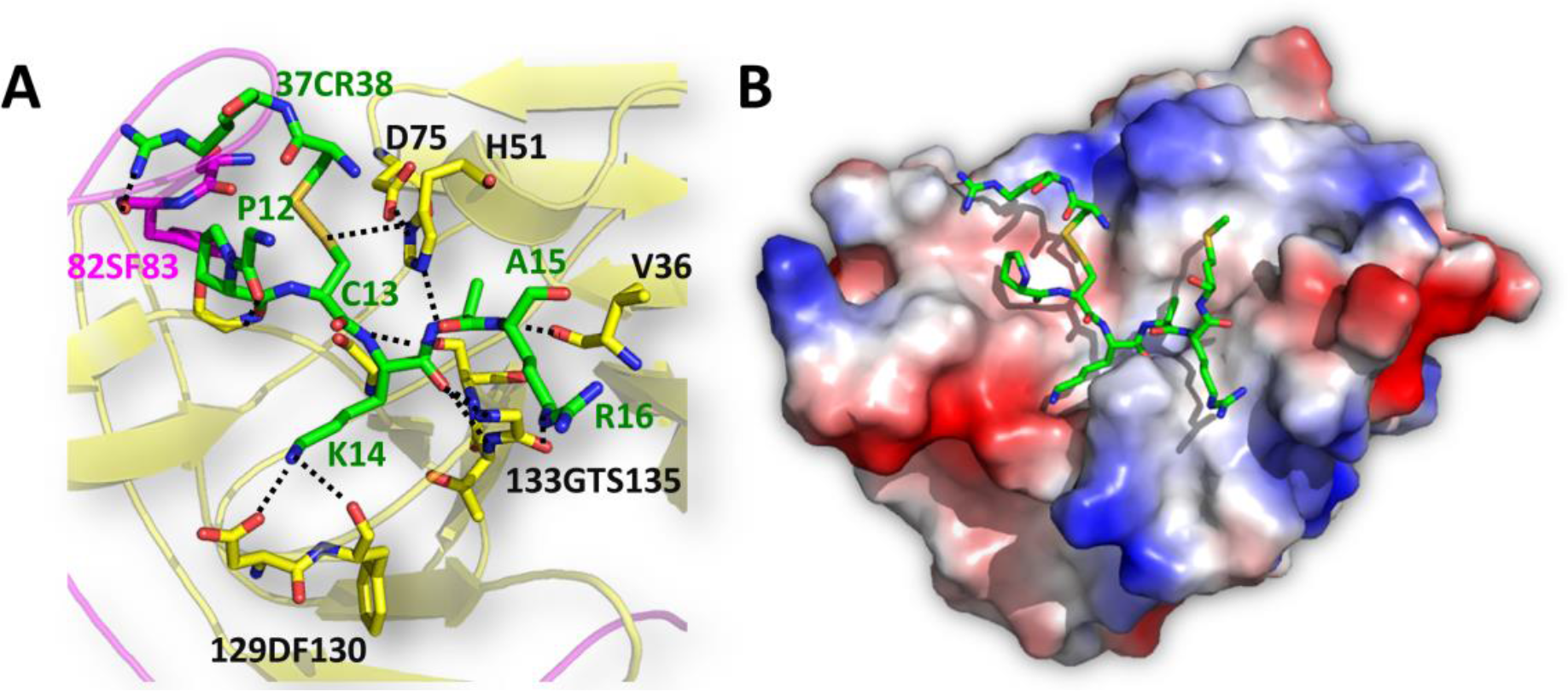
Binding of BPTI to NS3 protease is conserved among the different flavivirus protease structures. (A) Detailed interactions between BPTI and NS2B-NS3 protease. BPTI is presented in green, NS2B in magenta and NS3 in yellow. The interacting residues are shown as sticks. The residues numbers are labelled and color accordingly. (B) Electrostatic surface view of eNS2B_47_NS3 protease domain with bound BPTI residues in the pocket.

**S7 Fig.**
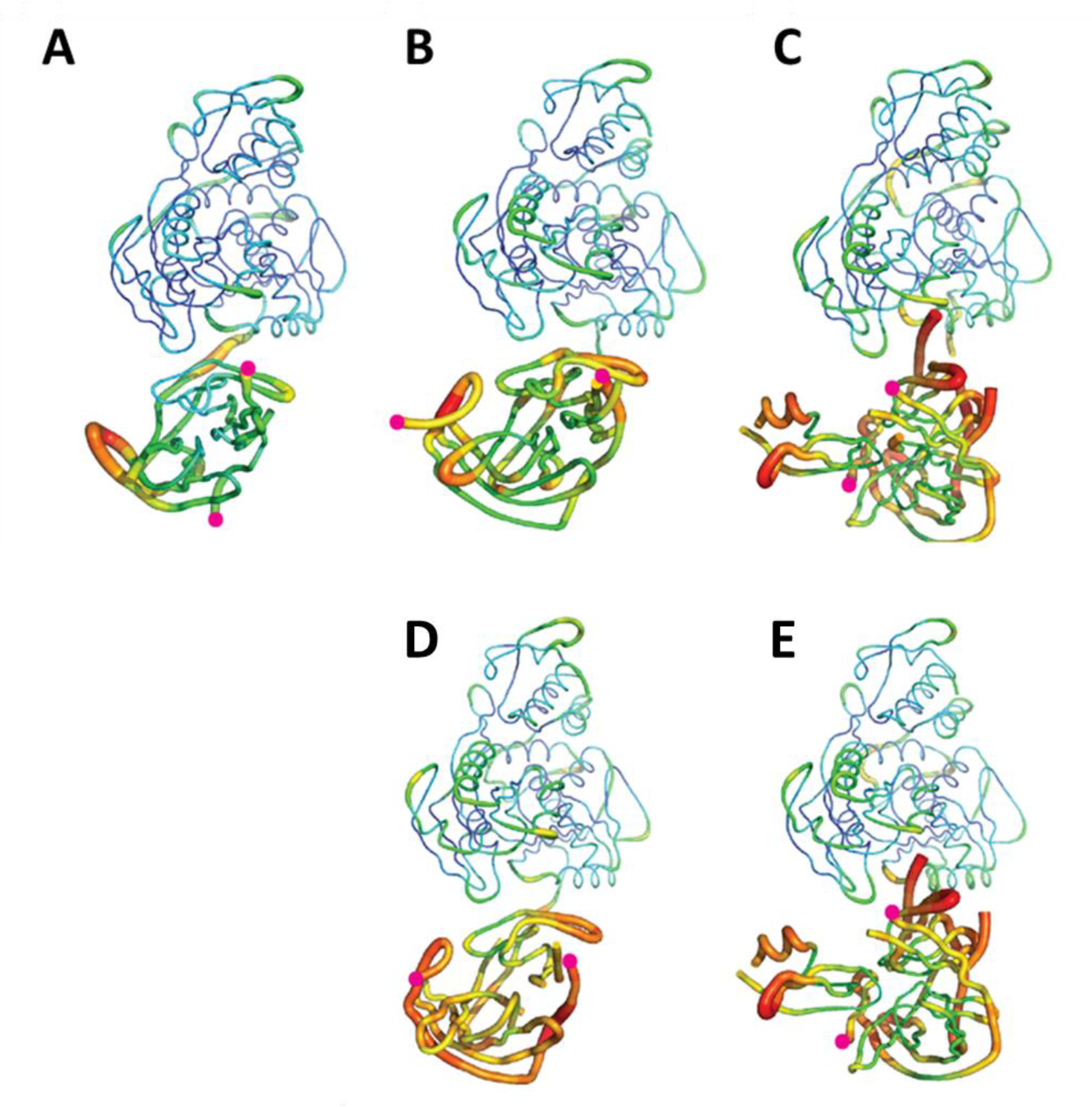
The b-factor putty representation of the full length crystal structures of NS2B_47_NS3 shows that protease domain is dynamic and changes conformations. (A)(B)(C) Full length crystal structures of gNS2B_47_NS3 (A) in open NS2B conformation (B) in closed NS2B conformation (C) in complex with BPTI. (D)(E) Full length crystal structure of unlinked eNS2B_47_NS3 (D) in closed NS2B conformation and (E) in complex with BPTI. Magenta dots represent the N- and C-terminus residues of NS2B.

**S1 Table.** Crystallisation conditions and unit cell dimensions of NS2B_47_NS3 constructs. All the crystal structures of NS2B_47_NS3 crystallised in three distinct unit cell dimensions (1) open conformation, (2) closed conformation and (3) closed in complex with BPTI. The glycine linker construct gNS2B_47_NS3 free enzyme structures are in both open and closed conformation without an inhibitor, where eNS2B_47_NS3 free enzyme structures are only captured in closed conformation regardless of the presence of inhibitor

| Construct name | Crystallisation conditions | Unit cell dimensions<br>(a b c,<br>α β γ) | Conformation |
| --- | --- | --- | --- |
| **gNS2B<sub>47</sub>NS3 S135A** | 0.1 M MES pH 6.4, 15% PEG<br>6000 | 52.78 **75.9** **86.81**<br>89.92 90.32 **93.07** | Open |
| **gNS2B<sub>47</sub>NS3 L30S F31S** | 0.1 M MES pH 6.4, 15% PEG | 52.92 **88.72** **81.30**<br>90 **93.08** 90 | Closed |
| **eNS2B<sub>47</sub>NS3 S135A** | 0.1M MES pH 6.4, 10% PEG<br>4000 | 52.77 88.3 81.11<br>90 **92.34** 90 | Closed |
| **eNS2B<sub>47</sub>NS3 L30S F31S** | 0.1 M MES pH 6.4. 10% PEG<br>6000 | 52.9 88.77 81.45<br>90 **93.85** 90 | Closed |
| **gNS2B<sub>47</sub>NS3 S135A + BPTI** | 0.1 MES pH 6.4, 12% PEG<br>6000 | 53.06 **85.63** **85.51**<br>90 **97.95** 90 | Closed, in complex with<br>BPTI |
| **eNS2B<sub>47</sub>NS3 L30S F31S +**<br>**BPTI** | 0.1 M MES pH 6.0, 12% PEG<br>4000 | 53.02 **87.493** **86.46**<br>90 **98.25** 90 | Closed, in complex with<br>BPTI |
| **bNS2B<sub>47</sub>NS3** | 0.1 M MES pH 6.0, 12% PEG<br>6000 | 52.65 **87.64** **80.12**<br>90 **91.94** 90 |  |

## References

1. Cao-Lormeau, V.M., et al., Guillain-Barre Syndrome outbreak associated with Zika virus infection in French Polynesia: a case-control study. Lancet, 2016. 387(10027): p. 1531–1539.

2. Brasil, P., et al., Zika Virus Infection in Pregnant Women in Rio de Janeiro. N Engl J Med, 2016. 375(24): p. 2321–2334.

3. Bell, B.P., C.A. Boyle, and L.R. Petersen, Preventing Zika Virus Infections in Pregnant Women: An Urgent Public Health Priority. Am J Public Health, 2016. 106(4): p. 589–90.

4. Chambers, T.J., et al., Flavivirus Genome Organization, Expression, and Replication. Annual Review of Microbiology, 1990. 44: p. 649–688.

5. Lindenbach, B.D. and C.M. Rice, Molecular biology of flaviviruses. Adv Virus Res, 2003. 59: p. 23–61.

6. Lescar, J., et al., Towards the design of antiviral inhibitors against flaviviruses: the case for the multifunctional NS3 protein from Dengue virus as a target. Antiviral Res, 2008. 80(2): p. 94–101.

7. Luo, D., S.G. Vasudevan, and J. Lescar, The flavivirus NS2B-NS3 protease-helicase as a target for antiviral drug development. Antiviral Res, 2015. 118: p. 148–158.

8. Ma, Y., et al., NS3 helicase domains involved in infectious intracellular hepatitis C virus particle assembly. J Virol, 2008. 82(15): p. 7624–39.

9. Falgout, B., et al., Both nonstructural proteins NS2B and NS3 are required for the proteolytic processing of dengue virus nonstructural proteins. J Virol, 1991. 65(5): p. 2467–75.

10. Chambers, T.J., A. Grakoui, and C.M. Rice, Processing of the yellow fever virus nonstructural polyprotein: a catalytically active NS3 proteinase domain and NS2B are required for cleavages at dibasic sites. J Virol, 1991. 65(11): p. 6042–50.

11. Noble, C.G., et al., Ligand-bound structures of the dengue virus protease reveal the active conformation. J Virol, 2012. 86(1): p. 438–46.

12. Erbel, P., et al., Structural basis for the activation of flaviviral NS3 proteases from dengue and West Nile virus. Nat Struct Mol Biol, 2006. 13(4): p. 372–3.

13. Incicco, J.J., et al., Steady-state NTPase activity of Dengue virus NS3: number of catalytic sites, nucleotide specificity and activation by ssRNA. PLoS One, 2013. 8(3): p. e58508.

14. Li, H., et al., The serine protease and RNA-stimulated nucleoside triphosphatase and RNA helicase functional domains of dengue virus type 2 NS3 converge within a region of 20 amino acids. J Virol, 1999. 73(4): p. 3108–16.

15. Benarroch, D., et al., The RNA helicase, nucleotide 5’-triphosphatase, and RNA 5’- triphosphatase activities of Dengue virus protein NS3 are Mg2+-dependent and require a functional Walker B motif in the helicase catalytic core. Virology, 2004. 328(2): p. 208–18.

16. Rupp, D. and R. Bartenschlager, Targets for antiviral therapy of hepatitis C. Semin Liver Dis, 2014. 34(1): p. 9–21.

17. Low, J.G., E.E. Ooi, and S.G. Vasudevan, Current Status of Dengue Therapeutics Research and Development. J Infect Dis, 2017. 215(suppl_2): p. S96–S102.

18. Kaptein, S.J. and J. Neyts, Towards antiviral therapies for treating dengue virus infections. Curr Opin Pharmacol, 2016. 30: p. 1–7.

19. Chambers, T.J., et al., Evidence That the N-Terminal Domain of Nonstructural Protein Ns3 from Yellow-Fever Virus Is a Serine Protease Responsible for Site-Specific Cleavages in the Viral Polyprotein. Proceedings of the National Academy of Sciences of the United States of America, 1990. 87(22): p. 8898–8902.

20. Leung, D., et al., Activity of recombinant dengue 2 virus NS3 protease in the presence of a truncated NS2B co-factor, small peptide substrates, and inhibitors. J Biol Chem, 2001. 276(49): p. 45762–71.

21. Noble, C.G. and P.Y. Shi, Structural biology of dengue virus enzymes: towards rational design of therapeutics. Antiviral Res, 2012. 96(2): p. 115–26.

22. Robin, G., et al., Structure of West Nile virus NS3 protease: ligand stabilization of the catalytic conformation. J Mol Biol, 2009. 385(5): p. 1568–77.

23. Zhang, Z., et al., Crystal structure of unlinked NS2B-NS3 protease from Zika virus. Science, 2016. 354(6319): p. 1597–1600.

24. Phoo, W.W., et al., Structure of the NS2B-NS3 protease from Zika virus after self-cleavage. Nat Commun, 2016. 7: p. 13410.

25. Chandramouli, S., et al., Serotype-specific structural differences in the protease-cofactor complexes of the dengue virus family. J Virol, 2010. 84(6): p. 3059–67.

26. Hammamy, M.Z., et al., Development and characterization of new peptidomimetic inhibitors of the West Nile virus NS2B-NS3 protease. ChemMedChem, 2013. 8(2): p. 231–41.

27. Aleshin, A.E., et al., Structural evidence for regulation and specificity of flaviviral proteases and evolution of the Flaviviridae fold. Protein Sci, 2007. 16(5): p. 795–806.

28. de la Cruz, L., et al., Binding of low molecular weight inhibitors promotes large conformational changes in the dengue virus NS2B-NS3 protease: fold analysis by pseudocontact shifts. J Am Chem Soc, 2011. 133(47): p. 19205–15.

29. de la Cruz, L., et al., Binding mode of the activity-modulating C-terminal segment of NS2B to NS3 in the dengue virus NS2B-NS3 protease. FEBS J, 2014. 281(6): p. 1517–33.

30. Su, X.C., et al., NMR analysis of the dynamic exchange of the NS2B cofactor between open and closed conformations of the West Nile virus NS2B-NS3 protease. PLoS Negl Trop Dis, 2009. 3(12): p. e561.

31. Su, X.C., et al., NMR study of complexes between low molecular mass inhibitors and the West Nile virus NS2B-NS3 protease. FEBS J, 2009. 276(15): p. 4244–55.

32. Kim, Y.M., et al., NMR analysis of a novel enzymatically active unlinked dengue NS2B-NS3 protease complex. J Biol Chem, 2013. 288(18): p. 12891–900.

33. Zhang, Z.Z., et al., Crystal structure of unlinked NS2B-NS3 protease from Zika virus. Science, 2016. 354(6319): p. 1597–1600.

34. Shannon, A.E., et al., Product release is rate-limiting for catalytic processing by the Dengue virus protease. Sci Rep, 2016. 6: p. 37539.

35. Luo, D., et al., Crystal structure of the NS3 protease-helicase from dengue virus. J Virol, 2008. 82(1): p. 173–83.

36. Xu, T., et al., Structure of the Dengue virus helicase/nucleoside triphosphatase catalytic domain at a resolution of 2.4 A. J Virol, 2005. 79(16): p. 10278–88.

37. Luo, D., et al., Flexibility between the protease and helicase domains of the dengue virus NS3 protein conferred by the linker region and its functional implications. J Biol Chem, 2010. 285(24): p. 18817–27.

38. Assenberg, R., et al., Crystal structure of a novel conformational state of the flavivirus NS3 protein: implications for polyprotein processing and viral replication. J Virol, 2009. 83(24): p. 12895–906.

39. Li, J., et al., Functional profiling of recombinant NS3 proteases from all four serotypes of dengue virus using tetrapeptide and octapeptide substrate libraries. J Biol Chem, 2005. 280(31): p. 28766–74.

40. Li, Y., et al., Structural characterization of the linked NS2B-NS3 protease of Zika virus. FEBS Lett, 2017. 591(15): p. 2338–2347.

41. Lin, K.H., et al., Dengue Virus NS2B/NS3 Protease Inhibitors Exploiting the Prime Side. J Virol, 2017. 91(10).

42. Clum, S., K.E. Ebner, and R. Padmanabhan, Cotranslational membrane insertion of the serine proteinase precursor NS2B-NS3(Pro) of dengue virus type 2 is required for efficient in vitro processing and is mediated through the hydrophobic regions of NS2B. J Biol Chem, 1997. 272(49): p. 30715–23.

43. Battye, T.G., et al., iMOSFLM: a new graphical interface for diffraction-image processing with MOSFLM. Acta Crystallogr D Biol Crystallogr, 2011. 67(Pt 4): p. 271–81.

44. Kabsch, W., Integration, scaling, space-group assignment and post-refinement. Acta Crystallogr D Biol Crystallogr, 2010. 66(Pt 2): p. 133–44.

45. Kabsch, W., Xds. Acta Crystallogr D Biol Crystallogr, 2010. 66(Pt 2): p. 125–32.

46. Evans, P.R., An introduction to data reduction: space-group determination, scaling and intensity statistics. Acta Crystallogr D Biol Crystallogr, 2011. 67(Pt 4): p. 282–92.

47. Evans, P.R. and G.N. Murshudov, How good are my data and what is the resolution? Acta Crystallogr D Biol Crystallogr, 2013. 69(Pt 7): p. 1204–14.

48. Potterton, E., et al., A graphical user interface to the CCP4 program suite. Acta Crystallogr D Biol Crystallogr, 2003. 59(Pt 7): p. 1131–7.

49. Winn, M.D., et al., Overview of the CCP4 suite and current developments. Acta Crystallogr D Biol Crystallogr, 2011. 67(Pt 4): p. 235–42.

50. Emsley, P. and K. Cowtan, Coot: model-building tools for molecular graphics. Acta Crystallographica Section D-Biological Crystallography, 2004. 60: p. 2126–2132.

51. Emsley, P., et al., Features and development of Coot. Acta Crystallographica Section D, 2010. 66(4): p. 486–501.

52. Adams, P.D., et al., PHENIX: a comprehensive Python-based system for macromolecular structure solution. Acta Crystallogr D Biol Crystallogr, 2010. 66(Pt 2): p. 213–21.

53. Afonine, P.V., et al., Towards automated crystallographic structure refinement with phenix.refine. Acta Crystallogr D Biol Crystallogr, 2012. 68(Pt 4): p. 352–67.

54. Headd, J.J., et al., Use of knowledge-based restraints in phenix.refine to improve macromolecular refinement at low resolution. Acta Crystallogr D Biol Crystallogr, 2012. 68(Pt 4): p. 381–90.

55. Hayward, S. and R.A. Lee, Improvements in the analysis of domain motions in proteins from conformational change: DynDom version 1.50. J Mol Graph Model, 2002. 21(3): p. 181–3.

56. Kotlovyi, V., W.L. Nichols, and L.F. Ten Eyck, Protein structural alignment for detection of maximally conserved regions. Biophys Chem, 2003. 105(2-3): p. 595–608.

57. DeLano, W.L., PyMOL molecular viewer: Updates and refinements. Abstracts of Papers of the American Chemical Society, 2009. 238.

58. DeLano, W.L., Use of PYMOL as a communications tool for molecular science. Abstracts of Papers of the American Chemical Society, 2004. 228: p. U313–U314.

59. Kiianitsa, K., J.A. Solinger, and W.D. Heyer, NADH-coupled microplate photometric assay for kinetic studies of ATP-hydrolyzing enzymes with low and high specific activities. Anal Biochem, 2003. 321(2): p. 266–71.

60. Lo, M.C., et al., Evaluation of fluorescence-based thermal shift assays for hit identification in drug discovery. Anal Biochem, 2004. 332(1): p. 153–9.

