## Supplementary material for "Crystal structures of full length DENV4 NS2B-NS3 reveal the dynamic interaction between NS2B and NS3": Suppl

### Crystal structures of full length NS3 from Dengue virus provide insights into dynamics of protease domain

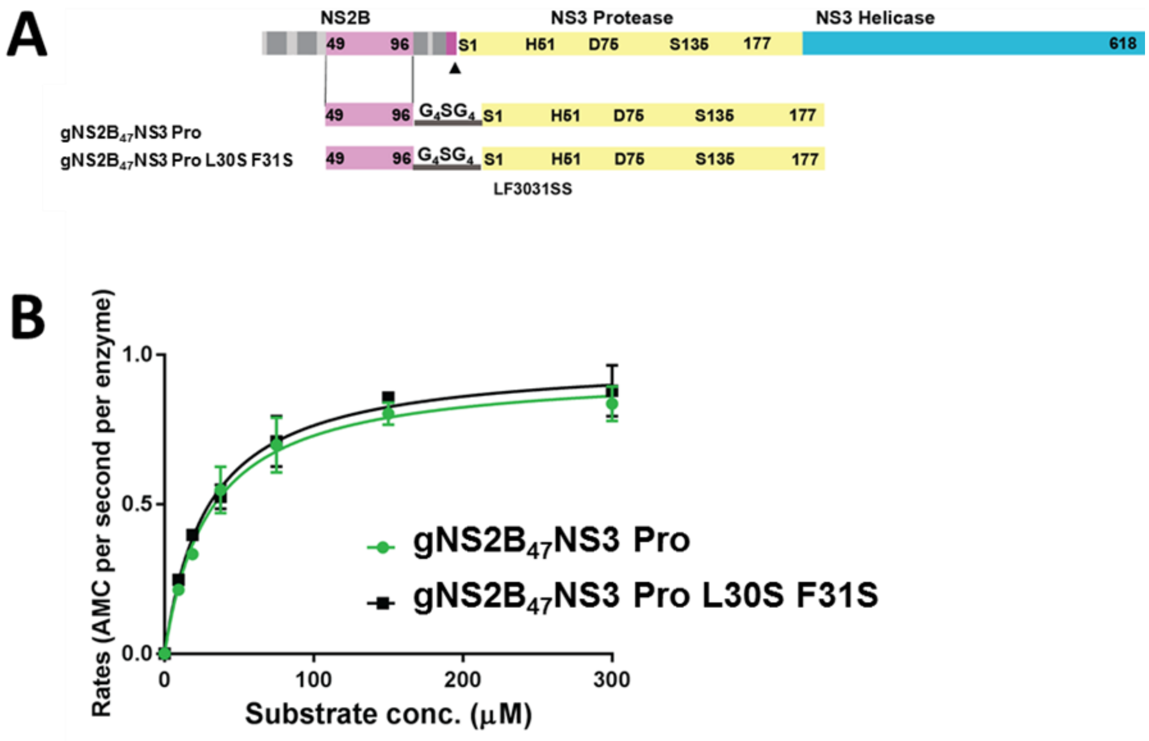


Supplementary Figure 1. The mutations L30S F31S do not interfere with protease activity. (A) Graphical representation of NS2B NS3 gene, construct design and mutations. NS2B in magenta, and NS3 protease in yellow and NS3 helicase in cyan. Protease and helicase are labelled and boundary residues are numbered. The catalytic residues, as well as mutated residues are labelled and numbered. (B) Enzymatic activity of $gNS2B_{47}NS3$ Pro and $gNS2B_{47}NS3$ Pro L30S F31S. Both enzymes have similar $k_{cat}$ and $K_m$ indicating that L30S F31S mutations do not interfere with enzymatic activity.

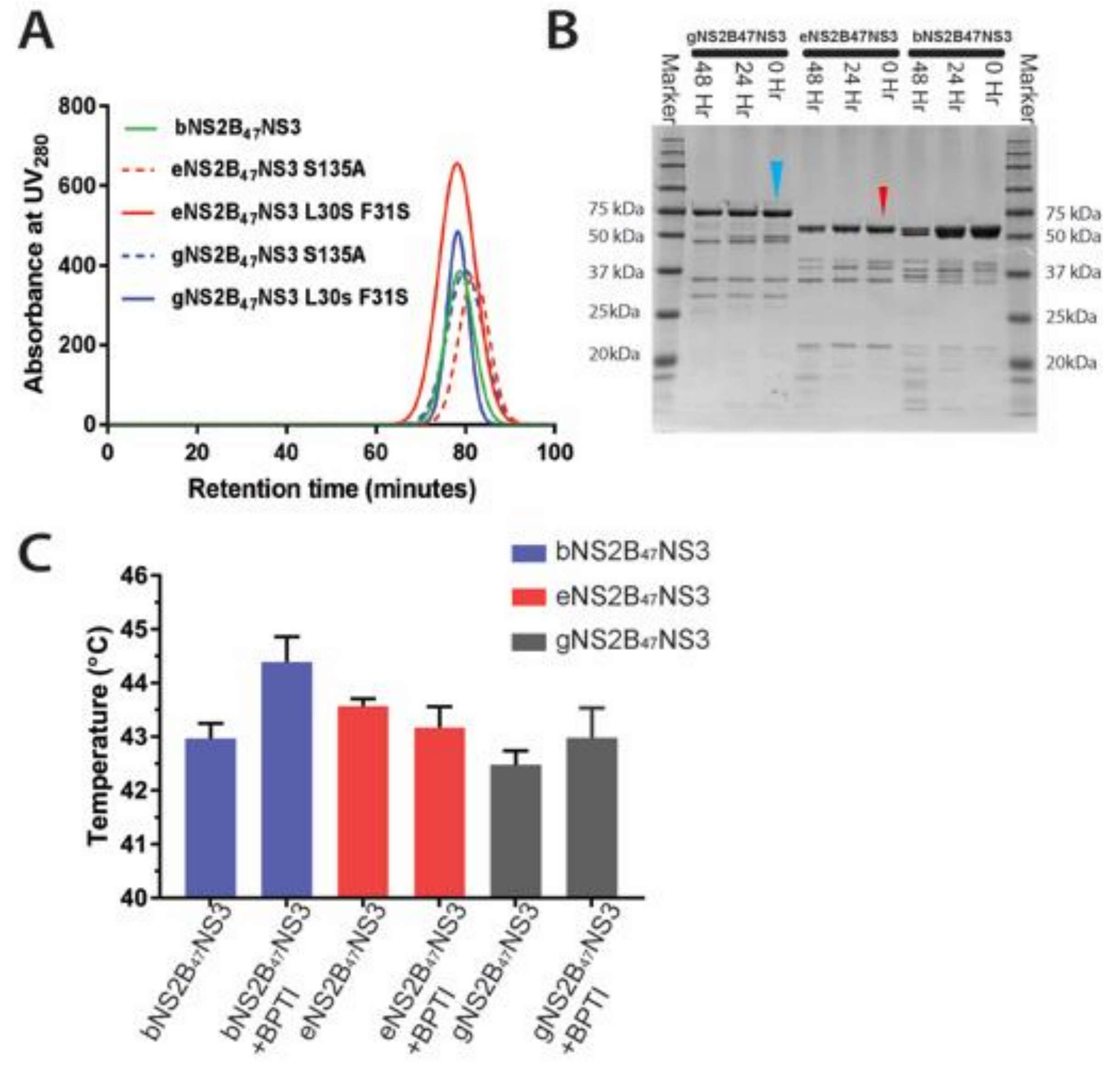


Supplementary Figure 2. Purification of gNS2B$_{47}$NS3, eNS2B$_{47}$NS3 and bNS2B$_{47}$NS3 showed monomeric proteins. (A) SEC chromatography profile of full length proteins showing that full length NS3 is monomeric. (B) SDS-PAGE analysis of full length NS3 auto proteolysis over 0hour, 24hour and 48 hours The gNS2B$_{47}$NS3 and eNS2B$_{47}$NS3 are indicated with blue and red arrows respectively.. (C) Melting temperatures of bNS2B$_{47}$NS3, eNS2B$_{47}$NS3 and gNS2B$_{47}$NS3 with/without BPTI. The gNS2B$_{47}$NS3 has the lowest $T_m$ indicating that gNS2B$_{47}$NS3 has lower stability compared to eNS2B$_{47}$NS3 and bNS2B$_{47}$NS3.

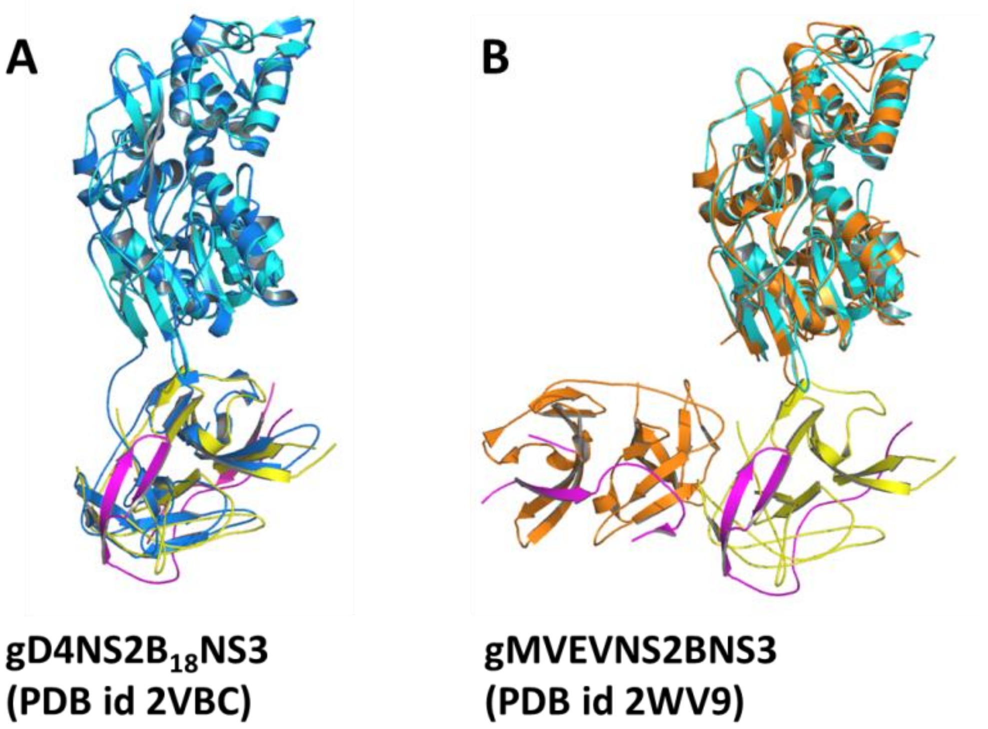


Supplementary Figure 3. Comparison between overall conformations of $NS2B_{47}NS3$ with previous structures. In both (A) and (B), $gNS2B_{47}NS3$ free enzyme structure in closed conformation was shown in cyan for helicase domain, yellow for protease and magenta for NS2B (A) Superposition of current full length $NS2B_{47}NS3$ with DENV4 full length NS3 structure with 18 residues from NS2B cofactor which is shown in blue for NS3 and magenta for NS2B (Luo et al PDB id 2VBC) (B) Superposition of current full length $NS2B_{47}NS3$ structure with MVEV $gNS2B_{47}NS3$ structure. The MVEV NS3 is shown in orange and NS2B in magenta.

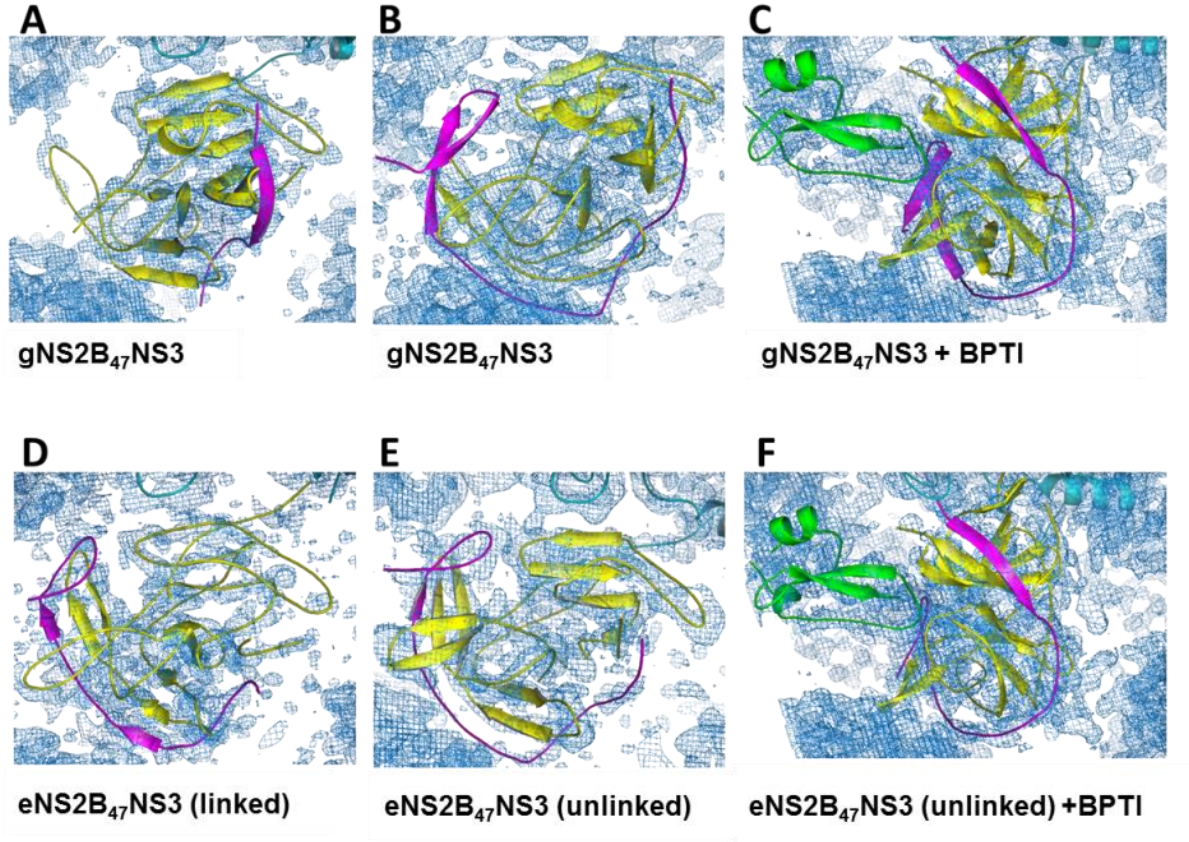


Supplementary Figure 5: The $2mF_o$-$F_c$ maps of $NS2B_{47}NS3$ structures display solvent shell around the electron density of protein. This indicates that the structure solutions for the protease domain are correct and refined. The NS2B is colored in magenta, NS3 protease in yellow and BPTI in green.

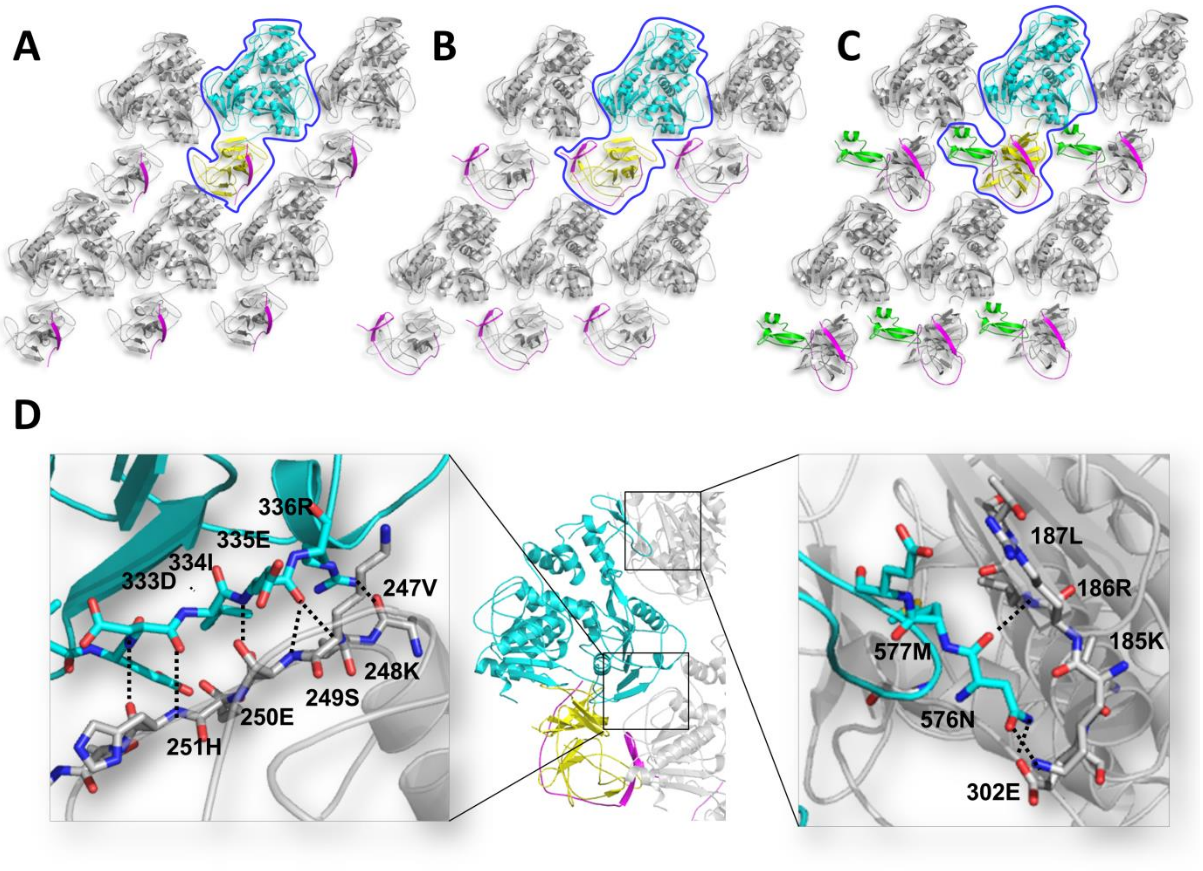


**Fig 4. Major crystal contacts in full length NS3 structures are formed by the helicase domain.** Here we display the three conformations of gNS2B$_{47}$NS3 full length structures (A) open NS2B conformation, (B) closed NS2B conformation, (C) enzyme in complex with BPTI along with its symmetry mates. NS3 helicase domain is shown in cyan and NS3 protease in yellow. The surrounding symmetry mates are shown in grey. The NS2B is shown magenta. (D) Detailed interactions of major crystal contacts. The residues that are interacting with the symmetry mates are presented with residue number. The protease domain does not interact with neighbouring molecules giving it the conformational freedom to adopt different orientations.

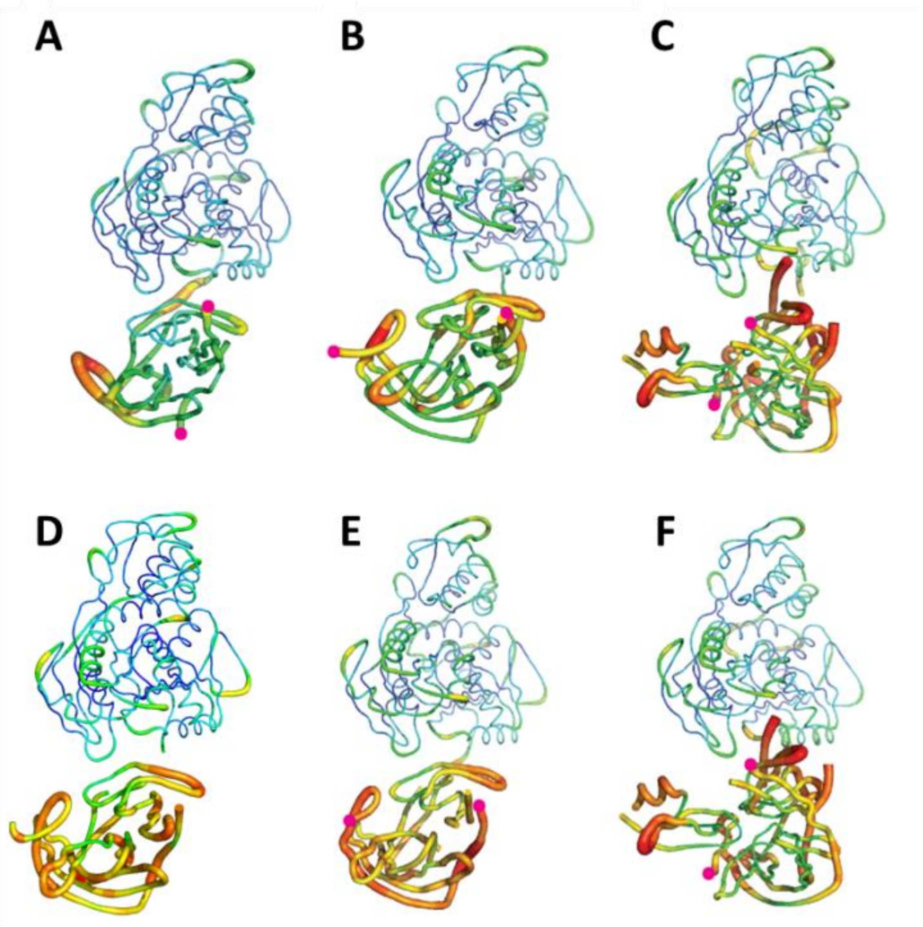


Supplementary Figure 8. The b-factor putty representation of the full length crystal structures of $NS2B_{47}NS3$ shows that protease domain is dynamic and changes conformations. (A)(B)(C) full length crystal structures of $gNS2B_{47}NS3$ (A) in open NS2B conformation (B) in closed NS2B conformation (C) in complex with BPTI. (D) full length crystal structure of linked $eNS2B_{47}NS3$ (E)(F) full length crystal structure of unlinked $eNS2B_{47}NS3$ (E) in closed NS2B conformation and (F) in complex with BPTI. Magenta dots represent the N- and C-terminus residues of NS2B.

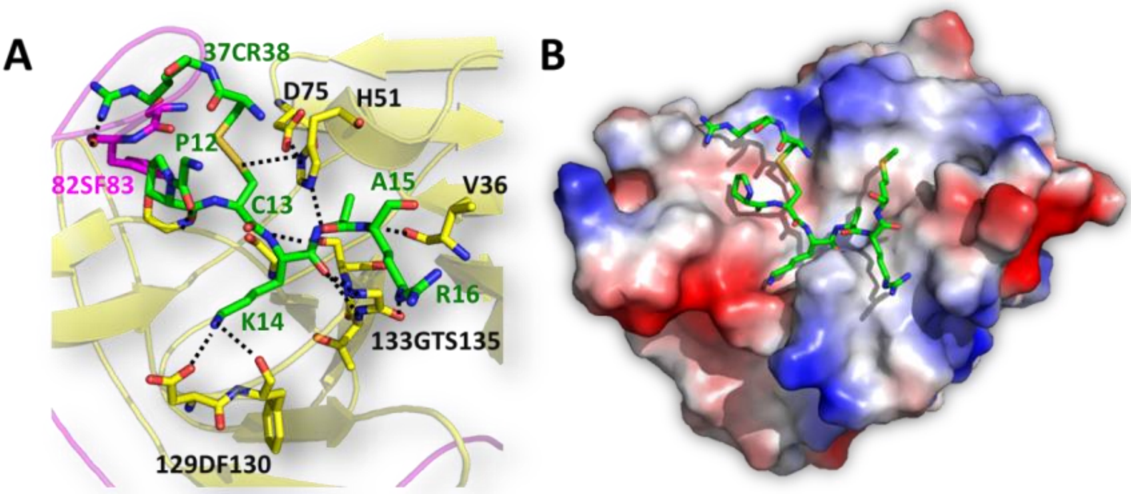


Supplementary Figure 7. Binding of BPTI to NS3 protease is conserved among the different flavivirus protease structures. (A) Detailed interactions between BPTI and NS2B-NS3 protease. BPTI is presented in green, NS2B in magenta and NS3 in yellow. The interacting residues are shown as sticks. The residues numbers are labelled and color accordingly. (B) Electrostatic surface view of eNS2B$_{47}$NS3 protease domain with bound BPTI residues in the pocket.

| Construct name | Crystallisation conditions | Unit cell dimensions (a b c, α β γ) | Conformation |
|---|---|---|---|
| **$gNS2B_{47}NS3$ S135A** | 0.1 M MES pH 6.4, 15% PEG 6000 | 52.78 **75.9 86.81** 89.92 90.32 **93.07** | Open |
| **$gNS2B_{47}NS3$ L30S F31S** | 0.1 M MES pH 6.4, 15% PEG | 52.92 **88.72 81.30** 90 **93.08** 90 | Closed |
| **Unlinked $eNS2B_{47}NS3$** | 0.1 M MES pH 6.4. 10% PEG 6000 | 52.9 88.77 81.45 90 **93.85** 90 | Closed |
| **$gNS2B_{47}NS3$ S135A + BPTI** | 0.1 MES pH 6.4, 12% PEG 6000 | 53.06 **85.63 85.51** 90 **97.95** 90 | Closed, in complex with BPTI |
| **Unlinked $eNS2B_{47}NS3$ + BPTI** | 0.1 M MES pH 6.0, 12% PEG 4000 | 53.02 **87.493 86.46** 90 **98.25** 90 | Closed, in complex with BPTI |
| **$bNS2B_{47}NS3$** | 0.1 M MES pH 6.0, 12% PEG 6000 | 52.65 **87.64 80.12** 90 **91.94** 90 | |

Supplementary Table 1. Crystallisation conditions and unit cell dimensions of $NS2B_{47}NS3$ constructs. All the crystal structures of $NS2B_{47}NS3$ crystallised in three distinct unit cell dimensions (1) open conformation, (2) closed conformation and (3) closed in complex with BPTI. The glycine linker construct $gNS2B_{47}NS3$ free enzyme structures are in both open and closed conformation without an inhibitor, where $eNS2B_{47}NS3$ free enzyme structures are only captured in closed conformation regardless of the presence of inhibitor
